# The dynamic evolution of panarthropod germ cell specification mechanisms

**DOI:** 10.1101/2025.08.05.668520

**Authors:** Jonchee A. Kao, Emily L. Rivard, Rishabh R. Kapoor, Cassandra G. Extavour

**Affiliations:** Department of Molecular and Cellular Biology, Harvard University, Cambridge MA, USA; Laboratory of Insect Physiology, School of Agriculture, Kyoto University, Kyoto, Japan; Department of Molecular Biology and Genetics, Cornell University, Ithaca NY, USA; Program for Systems, Synthetic and Quantitative Biology, Harvard University, Boston MA, USA; Department of Organismic and Evolutionary Biology, Harvard University, Cambridge MA, USA; Howard Hughes Medical Institute, Chevy Chase MD, USA

**Keywords:** germ plasm, pole cells, inductive signaling, BMP signaling pathway, cytoplasmic determinant

## Abstract

Germ cells enable the reproduction of an organism and the continuity of its lineage. Across animals, these crucial cells are segregated from the soma at different times and places and via distinct mechanisms. Understanding the evolution of germ cell specification across animals is complicated by the difficulty of making meaningful comparisons of embryonic development between diverse animal species. Here, we characterize germ cell specification in Panarthropoda, an ancient clade that encompasses massive animal biodiversity, within which we can conduct meaningful comparative embryology. We amass data from centuries of studies describing the timing and mechanisms of germ cell formation, and apply ancestral state reconstruction to these data to propose novel hypotheses about the trajectory of evolution in this process. Furthermore, we speculate about the mechanisms underlying these evolutionary dynamics by considering the relationships among germ cell specification, concurrent developmental processes and the germ line gene network. Collectively, this Review derives new insights from a rich historical database of embryological observations, offering broad implications for understanding the evolution of metazoan germ cells.

## Introduction

During animal development, the germ line comprises a collection of cells specified early in embryonic development that later give rise to the next generation via eggs or sperm. Whereas historical researchers relied primarily on observation of unperturbed embryos to understand germ cell specification and development (e.g. histological analysis of fixed, sectioned embryos; Bounoure, 1939; Wolff, 1962), we can now investigate these processes using functional genetic analysis. This allows us not only to identify germ cells with more certainty, but also to gain more insight into the molecular mechanisms of germ cell specification. In this Review, we have used “specification” to mean the process that imbues embryonic cells with the ultimate fate of generating eggs and sperm when the animal is sexually mature, and “differentiation” to mean the processes that make germ cells distinct from somatic cells in one or more of their morphology, behavior, or gene expression. Several hypotheses have been proposed to explain evolutionary transitions between the mechanisms underlying specification and differentiation across animals, including potential changes in cis-regulatory mechanisms, changes in evolvability or body plan diversity, or impacts on germ line mutation rate and ultimately fitness (Buss, 1982; Buss, 1983; Crother et al., 2007; Crother et al., 2016; Evans et al., 2014; Extavour, 2007; Johnson et al., 2011; Whittle and Extavour, 2016; Whittle and Extavour, 2017). However, such hypotheses are difficult to test because embryonic development of animals is too different across vast evolutionary time scales to be meaningfully compared. Moreover, there is disagreement in the field about whether existing well-studied mechanisms of specification (discussed below) are truly the only mechanisms, and whether they are discrete, mutually exclusive mechanisms. Furthermore, there are conflicting reports about the impact of germ cell specification mode on the evolution of genomes, body plans and fitness (Evans et al., 2014; Otto and Hastings, 1998; Queller, 2000; Whittle and Extavour, 2016; Whittle and Extavour, 2017). Here, we examine variation in germ cell specification across the most diverse clade of animals, the Panarthropoda, wherein the general features of embryogenesis can nevertheless be meaningfully compared. This clade displays striking diversity in the temporal and spatial origin of germ cells, as well as multiple transitions between specification mechanisms, offering an ideal system to study evolution and regulation of germ cell specification.

Here, we first review the existing observational data regarding embryonic germ line development across panarthropods. We are primarily interested in the first embryonic appearance of germ cells before their arrival in the gonad, during which time we refer to them as primordial germ cells (PGCs). We identify three categories of temporally distinct developmental context in which PGCs are first detectable during embryogenesis, and we analyze the phylogenetic distribution of these three categories. We then summarize perturbation-based experimental studies aimed at uncovering the mechanisms of germ cell specification. Finally, we discuss the implications of these data on our understanding of the evolution of germ cell specification in panarthropods, and more broadly across Metazoa.

### Identification of primordial germ cells

Historically, embryologists studied early development using histological preparations of sectioned embryos (His, 1868). With light microscopy, PGCs can be readily distinguished from surrounding somatic cells by their rounder and larger nuclei (Nieuwkoop and Sutasurya, 1979). The chromatin of germ cells often appears more diffuse than that of somatic cells, and the cytoplasm, especially cortical or perinuclear cytoplasm, may contain electron-dense, non-membrane bound granular material (Eddy, 1975). However, tracking these cells throughout development using fixed samples and morphological features alone is not always technically possible (reviewed in XXXX).

Since the late 20^th^ century, studies assessing gene expression patterns unique to or diagnostic of germ cells (henceforth referred to as molecular markers) have been instrumental in identifying germ cells at earlier stages than previously recognized. The most commonly used markers are the mRNA transcript or protein products (detectable by *in situ* hybridization or antibody staining respectively) of the genes *vasa*, *nanos*, *tudor* and *piwi* (Ewen-Campen et al., 2010). The use of molecular markers can enable germ cell identification at earlier developmental stages, since some germ cells express the genes that encode these molecules before their morphological differentiation becomes unambiguously detectable (e.g. Ewen-Campen et al., 2013a). Additionally, these markers facilitate tracking of germ cells over the course of development. However, using molecular markers is a biased approach, as it relies on previous knowledge of germ line gene expression from other species. Furthermore, interpretation of molecular marker data can be complicated by pleiotropy, as the genes that encode these molecules may also be expressed by somatic cells (e.g. Yajima and Wessel, 2011).

#### Germ cell formation in panarthropods

In our review of the embryological literature from 1860 to the present, we identified three categories of PGC specification in panarthropods, based on the timing of specification during embryogenesis (Fig. 1). We describe these categories below, along with relevant details of embryogenesis for context.

**Figure 1.**
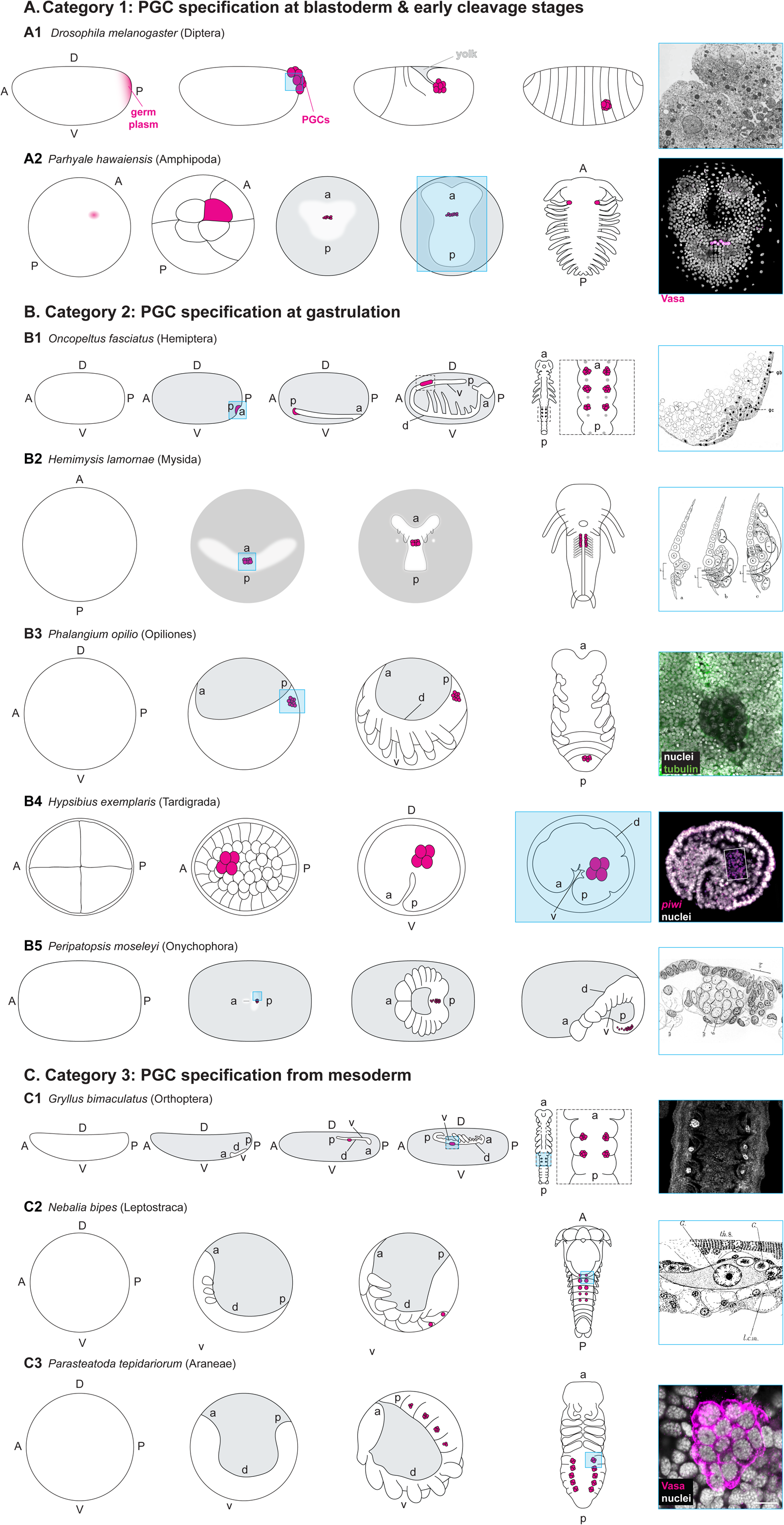
Primordial germ cell (PGC) origin in the context of panarthropod embryogenesis. In all schematics, A/a, anterior; P/p, posterior; D/d, dorsal; V/v, ventral.lower case letters are shown in cases where egg and embryonic axes are not the same, and/or indicate embryonic axes in embryos that do not occupy the entire egg volume; if D/V is not indicated, schematics show ventral view; magenta, approximate location of germ plasm or PGCs; gray, yolk or extraembryonic tissue; blue box in schematic, stage/location where germ cells are detectable in the micrographs and drawings on the right. (**A**) Description of Category 1: PGC specification at early cleavage and blastoderm stages. (**A1**) In the fruit fly *Drosophila melanogaster*, germ plasm accumulates at the posterior of syncytial embryos and is inherited by pole cells that migrate to the ultimate site of the embryonic gonad (Illmensee and Mahowald, 1974). Right: TEM micrograph displaying pole cells budding in *D. melanogaster* embryos. The pole cell nuclei (N) have characteristically diffuse chromatin, and polar granules (P) are distributed throughout the cytoplasm. Image reproduced with permission from (Mahowald, 1968). M: mitochondria; N: nuclei; P: polar granules. (**A2**) In the amphipod *Parhyale hawaiensis*, germ plasm accumulates in a region of the embryo called the RNA-containing body, which is inherited by the *g* micromere at the 8-cell stage (Extavour, 2005; Gupta and Extavour, 2013). All descendants of *g* are PGCs. Right: PGCs visualized by immunohistochemistry with a cross-reactive antibody raised against recombinant Vasa protein from both *D. melanogaster* and the grasshopper *Schistocerca americana* (Chang et al., 2002). (**B**) Description of Category 2: PGC specification at gastrulation. (**B1**) In the milkweed bug *Oncopeltus fasciatus*, PGCs first appear in the ventral posterior blastoderm in an invaginating structure called the posterior pit (Ewen-Campen et al., 2013a). PGCs remain at the posterior tip of the germ band as it invaginates into the yolk. The PGCs migrate along the germ band toward the anterior to segments A4–A6. Right: Hand drawing of a light micrograph of a sectioned late blastoderm stage *O. fasciatus* embryo showing germ cells in the posterior pit. Image reproduced with permission from (Butt, 1949). gb, germ band; gc, germ cell. (**B2**) In the mysid *Hemimysis lamornæ,* PGCs differentiate in the center of the blastopore area and are internalized at gastrulation (Manton, 1928). Right: Drawing of sagittal view of an *H. lamornæ* embryo at gastrulation. Image reproduced with permission from (Manton, 1928). B, blastopore area; E, ectodermal teloblast; en, endoderm cell; G, genital rudiment; M, mesodermal teloblast; m, head mesoderm band cell. (**B3**) In the harvestman *Phalangium opilio*, PGCs are first identifiable at the posterior end of the germ band soon after the germ band forms (Faussek, 1888; Gainett et al., 2022). PGCs remain at the posterior end of the germ band at least until the fourth opisthosomal segment is formed, after which the germ cells can be found in that segment. Right: Chromatin (visualized with the nucleic acid dye Hoechst 33342) is more diffuse in PGCs than in surrounding cells. Image kindly provided by Prashant Sharma (University of Wisconsin, Madison). Green, alpha-tubulin. (**B4**) In the tardigrade *Hypsibius exemplaris*, PGCs are first identifiable in the 32-cell stage and are the first cells to be internalized during gastrulation. Right: PGCs visualized by *in situ* hybridization against *He*-*vasa*. Image kindly provided by Kira Heikes (Duke University). (**B5**) In the velvet worm *Peripatopsis moseleyi*, PGCs are first identifiable near the blastopore at the beginning of gastrulation. As the embryo elongates and more segments form, the PGCs migrate to more anterior segments. Right: Hand drawing of a light micrograph of a *P. moseleyi* embryo showing primordial germ cells soon after their internalization during gastrulation, distinguished by their lighter-staining nuclei and cytoplasm. Image reproduced with permission from (Manton, 1949). G, germ cells; en, endoderm; b.a., blastoporal area. (**C**) Description of Category 3: PGC specification from mesoderm. (**C1**) In the cricket *Gryllus bimaculatus*, PGCs are first detectable among the mesoderm in segments A2–A4 at embryonic stage 6, after most abdominal segments have been specified (Ewen-Campen et al., 2013b). Right: PGCs visualized by immunohistochemistry with an antibody against *Gb*-Piwi. (**C2**) In the leptostracan *Nebalia bipes*, PGCs are first identifiable in the mesoderm in the thorax and abdomen during the extended germ band stage (Manton, 1934). Right: hand drawing of a light micrograph of a sectioned *N. bipes* embryo showing PGCs among the mesoderm in the extended germ band, distinguished by their lighter-staining cytoplasm and larger nuclei. Image reproduced with permission from (Manton, 1934). G, genital rudiment; l.c.m., circular muscle round liver lobe; th.8, eighth thoracic segment. (**C3**) In the house spider *Parasteatoda tepidariorum*, PGCs are first identifiable in segmental clusters in the opisthosomal mesoderm after all segments have formed (Schwager et al., 2015). Right: PGCs visualized by immunohistochemistry using an antibody raised against *Pt*-Vasa.

### Category 1: PGC specification at early cleavage and blastoderm stages

After fertilization, embryogenesis begins with cleavage, defined as a series of mitotic divisions of the zygote (Gilbert, 1997). In most arthropods, cleavage leads to the formation of the *blastoderm*, a relatively uniform layer of cells at the egg cortex, surrounding a central yolk mass populated by a small number of yolk cells (Anderson, 1973; Scholtz and Wolff, 2013). The first category of PGC specification in panarthropods occurs during or before the formation of the blastoderm (Fig. 1A).

#### In insects

In most insects, the blastoderm is initially a syncytium, which is a single cell containing multiple nuclei generated by incomplete cleavage division cycles that do not include cytokinesis (Anderson, 1973; Johannsen and Butt, 1941). In some insects, the PGCs are the first cells to form, appearing at the posterior pole before the rest of the blastoderm is cellularized (Fig. 1A1), and therefore often referred to as pole cells. While pole cells are most commonly seen in species of Diptera (flies, midges and mosquitoes), Coleoptera (beetles), and Hymenoptera (bees, ants and wasps), they have also been reported in Thysanoptera (thrips) and Dermaptera (earwigs) (references in Table S1).

Pole cells were first reported in a species of the midge *Chironomus* by Weismann (1863), although Metschnikow (1866) was the first to report their migration into the gonad. In many insects, the cytoplasm at the posterior pole of the embryo is morphologically distinct from that of the rest of the embryo, often with visible granules (Hegner, 1914; Zissler, 1992). The close association of this granular cytoplasm with pole cells led researchers to speculate that the granular material contains germ line determinants (e.g. Hegner, 1909a; Zissler, 1992). It was long believed that pole cells gave rise to somatic cell types in addition to the germ cells (Anderson, 1962a; Anderson, 1962b; Poulson, 1947; Poulson, 1950). However, careful lineage tracing in the fruit fly *Drosophila melanogaster* showed that all pole cells are determined as germ cells, and any that become lost during their migration die (Underwood et al., 1980).

While the function of pole cells as PGCs appears conserved across insects, the specific details of their formation can vary considerably across species. For example, pole cells differ in which cleavage cycle they cellularize: they form during the ninth mitotic cycle of *D. melanogaster* (Huettner, 1923) and during the sixth mitotic cycle in the parasitic wasp *Nasonia vitripennis* (Buchta et al., 2013). To our knowledge, the earliest observed formation of pole cells is during the third cleavage cycle in the midges *Miastor metraloas* and *Chironomus sp*. (Hegner, 1914; Weismann, 1863). The number of pole cells formed is also variable: a single pole cell forms in *Chironomus* embryos (Hegner, 1914; Weismann, 1863), whereas four appear in the fruit fly *Bactrocera tryoni* (Anderson, 1962a) and sixteen pole cells form in the beetle *Calligrapha multipunctata* (Hegner, 1909b).

After the pole cells form, they enter the egg and migrate to the location of the future embryonic gonad, which is quite far from the posterior pole (Santos and Lehmann, 2004). Several different migration pathways have been described. In *D. melanogaster* and many other Diptera, the pole cells are carried into the egg during posterior midgut invagination, then migrate through the epithelium of the posterior midgut to enter the abdominal mesoderm and associate with the developing somatic gonad (Metschnikow, 1866; Sonnenblick, 1941). In Hymenoptera, pole cells are internalized with the rest of the mesoderm during gastrulation (Gatenby, 1917). Yet another variation is observed in Chrysomelidae (leaf beetles) (Hegner, 1909a), where germ cells are internalized with the rest of the germ band as the extraembryonic membranes fold over the germ band.

#### In other arthropods

PGCs are also specified early in Collembola (springtails), a clade of hexapods that are sister to insects (Du et al., 2024). Embryogenesis in these animals begins with a period of holoblastic cleavage, and certain blastomeres on the basal (yolk) side of the blastoderm can be recognized as likely PGCs very early in development (Jura, 1967; Klag, 1982). The cytoplasm of these cells contains granular material visible by light and electron microscopy (Klag, 1982), resembling the cytoplasm containing germ line determinants observed in insects (Hegner, 1914; Zissler, 1992). Rather than maintaining contact with the germ band as in insects, collembolan germ cells migrate into the center of the yolk after their specification until eventually migrating to associate with the abdominal mesoderm, from which the embryonic gonad develops (Jura, 1967).

Among non-hexapod Pancrustacea, embryos of Copepoda (copepods), Amphipoda (beach hoppers), Decapoda (crabs, lobsters and shrimp), and some Cladocera (water fleas) undergo stereotypical *holoblastic cleavage*, allowing precise tracking of cell lineages and the identification of a specific cleavage blastomere as the progenitor of PGCs (Gerberding et al., 2002; Hertzler, 2005; Kühn, 1913; McClendon, 1907). In amphipods and decapods, a putative germ plasm has been identified in the freshly laid egg, termed the RNA-containing body (Gupta and Extavour, 2013) or intracellular body (Biffis et al., 2009; Chen et al., 2021). During cleavage, this region of cytoplasm is inherited asymmetrically by the blastomeres that give rise to the PGCs (Gupta and Extavour, 2013) (Fig. 1A2).

In chelicerates (spiders, scorpions, ticks and mites), candidate PGCs are first detectable with molecular markers in the yolk of blastoderm-stage embryos in the mite *Tetranychus urticae* (Dearden et al., 2003). As the germ band forms, the PGCs associate with the posterior germ band near the primordia of the fourth walking legs (Dearden et al., 2003). We are not aware of any other chelicerate or myriapod (centipedes and millipedes) species in which PGC specification has been proposed to occur before gastrulation.

### Category 2: PGC specification at gastrulation

After formation of the blastoderm, a portion of the blastoderm is specified as the presumptive embryonic region, variably referred to as the *germ band*, *germ rudiment*, *embryonic rudiment* or *germ disc* depending on its initial form (Anderson, 1973; Johannsen and Butt, 1941). The rest of the blastoderm differentiates into one or more extraembryonic membranes (Johannsen and Butt, 1941). Soon after this specification, *gastrulation* occurs, whereby some of the blastoderm cells are internalized and become the mesoderm (Stern, 2004). In insects, mesoderm internalization primarily occurs along the ventral midline of the germ band (Leptin, 2004; Roth, 2004). In other arthropods, mesoderm may be internalized through a *blastopore* near the germ rudiment (Chaw et al., 2007; Gerberding and Patel, 2004), or by epiboly (Edgar et al., 2015). The second category of PGC differentiation occurs at or very close to the time of gastrulation (Fig. 1B).

#### In insects

Among insects, PGC specification during gastrulation is the most commonly reported timing in members of Hemiptera (true bugs), Psocodea (lice) and Lepidoptera (moths and butterflies), although it is also reported in species of Dermaptera, Blattodea (cockroaches), Hymenoptera, Megaloptera (alderflies), Strepsiptera (twisted-wing flies), Coleoptera, Mecoptera (scorpion flies) and Diptera. Three different spatial patterns of germ cell specification have been reported around the time of gastrulation: before germ band formation, at the posterior of the germ band after its formation, or at the ventral midline of the blastoderm just before gastrulation. In Hemiptera, Psocodea and Tenebrionidae (darkling beetles; Coleoptera), germ cells are first detectable at the ventral posterior region of the cellular blastoderm in an invaginating structure called the *posterior pit* (Fig. 1B1) (Butt, 1949; Goss, 1952; Mellanby, 1935; Schröder, 2006; Shinji, 1919). These cells appear to form on the side of the blastoderm epithelium facing the yolk, rather than on the outside of the blastoderm, such as the pole cells described in Category 1. In Hemiptera and Psocodea, the invagination continues into the egg towards the anterior, forming a definitive germ band with germ cells at its posterior tip (Fig. 1B1). Germ cells in tenebrionid beetles move further inside the embryo once the amnion (ventral extraembryonic membrane) encloses the entire ventral surface of the germ band, similar to the internalization seen in coleopteran species that specify their germ cells as pole cells (e.g. the weevils *Callosobruchus maculatus* (Brauer, 1925; Brauer, 1949; Quan, 2018) and *Acanthoscelides obtectus* (Jung, 1966; Lynch et al., 2011)). The newly formed germ cells then migrate along the dorsal side of the germ band toward the anterior, eventually settling into the somatic gonad in the abdominal mesoderm.

In several species that appear to specify germ cells around the time of gastrulation, PGCs were first identified at the posterior tip of the germ band after its detachment from the blastoderm [*e.g.*, *Periplaneta orientalis* (Blattodea) (Heymons, 1895), *Sialis mitsuhashii* (Megaloptera) (Suzuki et al., 1981), *Nematus ribesii* (Hymenoptera) (Singh, 1967), and Mecoptera (Suzuki, 1990)]. Early studies of tenebrionid beetles described this pattern of germ cell differentiation as well (Rempel and Church, 1969; Ullmann, 1964). However, later work using molecular markers suggests that PGCs in at least one such beetle, *Tribolium castaneum*, appear earlier, during the cellular blastoderm stage in a posterior pit much like the Hemiptera (Schröder, 2006). This raises the possibility that in some other tenebrionid species, PGCs may likewise differentiate from the blastoderm, with the reported later appearance in the germ band simply reflecting limitations in detection methods.

In contrast to the posterior pit, germ cells differentiate along the ventral midline of the blastoderm in Lepidoptera, separating from the blastoderm before gastrulation to lie on the dorsal side of the presumptive germ band in the presumptive abdominal region (Ando and Tanaka, 1980; Miya, 1958a; Presser and Rutschky, 1957; Tanaka, 1987; Woodworth, 1889). Unlike when germ cells form at the posterior pit or at the posterior tip of the germ band, in Lepidoptera there does not appear to be long-range germ cell migration, since the germ cells differentiate in the same abdominal location as the gonad.

#### In other panarthropods

Among non-insect Pancrustacea, PGCs are the first cells to be internalized from the blastopore lip in Mysida (opossum shrimp) (Manton, 1928), Anomopoda (water fleas) (Cannon, 1921), and the copepod orders Siphonostomatoida (sea lice) (McClendon, 1907), Harpacticoida (Witschi, 1934) and Cyclopoida (Schimkewitsch, 1896). In Isopoda (woodlice and pill bugs), PGCs differentiate from the mesendoderm soon after internalization (Fig. 1B2) (Goodrich, 1939; Needham, 1942; Strömberg, 1965; Wolff, 2009). Thus, PGC differentiation seems to be tied to gastrulation in these crustacean orders.

In chelicerates, PGCs have been identified based on morphological criteria at the blastopore in Opiliones (harvestmen) (Fig. 1B3) (Faussek, 1888; Faussek, 1892; Holm, 1947) and Scorpiones (Brauer, 1894), and a recent study using molecular markers supported this interpretation (Gainett et al., 2022). Putative PGCs have also been identified at the blastopore in the centipedes *Scolopendra cingulata* and *Strigamia maritima* (Myriapoda) (Green and Akam, 2014; Heymons, 1901).

In Tardigrada (water bears), germ cells are the first cells to enter the blastopore during gastrulation, based on data from both morphological characteristics (Hejnol and Schnabel, 2005) and molecular markers (Heikes et al., 2023) (Fig. 1B4). Lineage tracing in these embryos suggests that the germ cells are derived from both blastomeres of the two-cell embryo (Gabriel et al., 2007; Hejnol and Schnabel, 2005).

Finally, onychophoran (velvet worm) germ cells have been described only with classic histological methods, to our knowledge (Fig. 1B5). They emerge either at the blastopore lip during gastrulation (Manton, 1949) or at the posterior end of the germ band during elongation (Manton, 1949; Mayer and Tait, 2009; Sedgwick, 1887).

### Category 3: PGC specification from mesoderm

After gastrulation, the germ band elongates and becomes segmented, and the mesoderm becomes organized into pairs of somites in each segment. In most living arthropods, each somite forms a small *coelom* enclosing a *coelomic cavity* (Anderson, 1973; Johannsen and Butt, 1941). The coelomic cavities are eventually lost as the mesoderm differentiates into different cell types and reorganizes into the primordia of the internal organ systems, including the somatic gonad (Anderson, 1973; Johannsen and Butt, 1941). The final category of PGC differentiation timing encompasses species in which PGCs differentiate from the mesoderm at this comparatively late stage of embryogenesis (Fig. 1C).

#### In insects

In insects, PGC specification from the mesoderm has been reported in members of Archaeognatha (jumping bristletails), Zygentoma (silverfish and firebrats), Orthoptera (crickets and grasshoppers) (Fig. 1C1), Mantodea (praying mantises), Blattodea, Trichoptera (caddisflies), Coleoptera and Lepidoptera. PGCs may differentiate before formation of the coelomic cavities [e.g., Orthoptera (Ewen-Campen et al., 2013b), Mantodea (Görg, 1959)], after formation of the coelomic cavities [e.g., Archaeognatha (Larink, 1969), Zygentoma (Woodland, 1957)], or after the coelomic cavities are no longer recognizable [e.g., Trichoptera (Miyakawa, 1974), *Odontotaenius disjunctus* (Coleoptera) (Krause, 1947), *Apis mellifera* (Hymenoptera) (Dearden, 2006; Nelson, 1915)]. In this developmental timing category, PGCs are usually specified in the same abdominal segments where the somatic gonad forms, so there is no long-range movement of PGCs after their specification.

#### In other arthropods

Differentiation of PGCs from the mesoderm has also been described in Leptostraca (mud shrimp) (Figure 1C2) (Manton, 1934), Anaspidacea (mountain shrimp) (Hickman, 1936) and Branchiopoda (fairy, clam and tadpole shrimp, and water fleas) (Anderson, 1967; Mitsumoto and Makioka, 2002; Sudler, 1899). However, data for stages of crustacean development late enough to include differentiation of the trunk mesoderm from which primordial germ cells might arise are sparse. This relative scarcity of early embryonic descriptions may be in part because many crustaceans hatch as a nauplius, comprising only the head segments (Scholtz, 2000). In these animals, germ cells might not be specified until later larval stages, after formation of the trunk. Many studies of naupliar and larval development in crustaceans are focused on organogenesis of the gonad as a whole, without specific reference to the origin of PGCs (Cannon, 1924; Ikuta and Makioka, 1994; Mitsumoto and Makioka, 2002).

Classical studies on myriapods suggest a late differentiation of PGCs from the mesoderm in three of the four myriapod classes [Pauropoda (Tiegs, 1947), Symphyla (pseudocentipedes) (Tiegs, 1940) and Diplopoda (millipedes) (Dohle, 1964)], although only one species was reported from each class in these studies. Germ cells in these myriapod classes have not been investigated with molecular markers to our knowledge.

Among chelicerates, PGCs differentiate from the mesoderm of the opisthosomal segments in spiders (Fig. 1C3) (Gough, 1902; Kautzsch, 1910; Schimkewitsch, 1903), consistent with data from molecular markers are (Schwager et al., 2015). In pycnogonids (sea spiders), PGCs are first detectable in the mesoderm of the prosoma (Alexeeva and Tamberg, 2021; Alexeeva et al., 2018; Miyazaki and Makioka, 2012).

### Summary: evolutionary trends from late to early germ cell specification

In his monograph on embryogenesis in Dermaptera and Orthoptera, Richard Heymons (1895) proposed that ‘the formation of sex cells at the rear end of the embryonic primordium, which has now been clearly demonstrated for representatives of very different groups, is probably generalizable for all insects’. Rather than making broad statements such as this, we summarize the data regarding the timing of germ cell specification (Figs. 2–3; Table S1) in the context of currently accepted panarthropod phylogeny (Ballesteros et al., 2022; Fernández et al., 2018; Giribet and Edgecombe, 2019; Misof et al., 2014; Schwentner et al., 2017). This shows that in euarthropod orders that are close to the base of the tree, PGCs tend to be first observed later in embryogenesis (discussed above as Category 3). This pattern suggests that the timing of first germ cell appearance evolved to be earlier in development in multiple lineages independently, as seen in Dermaptera, Condylognatha (true bugs and thrips) and Holometabola (undergoing complete metamorphosis) within insects; Copepoda, Malacostraca (the largest class of crustaceans) and Cladocera within non-hexapod Pancrustacea; Chilopoda (centipedes) within myriapods; and Opiliones, scorpions and mites within chelicerates. These data suggest the possibility that differentiation from the mesoderm is the ancestral mode of PGC specification in Euarthropoda.

**Figure 2.**
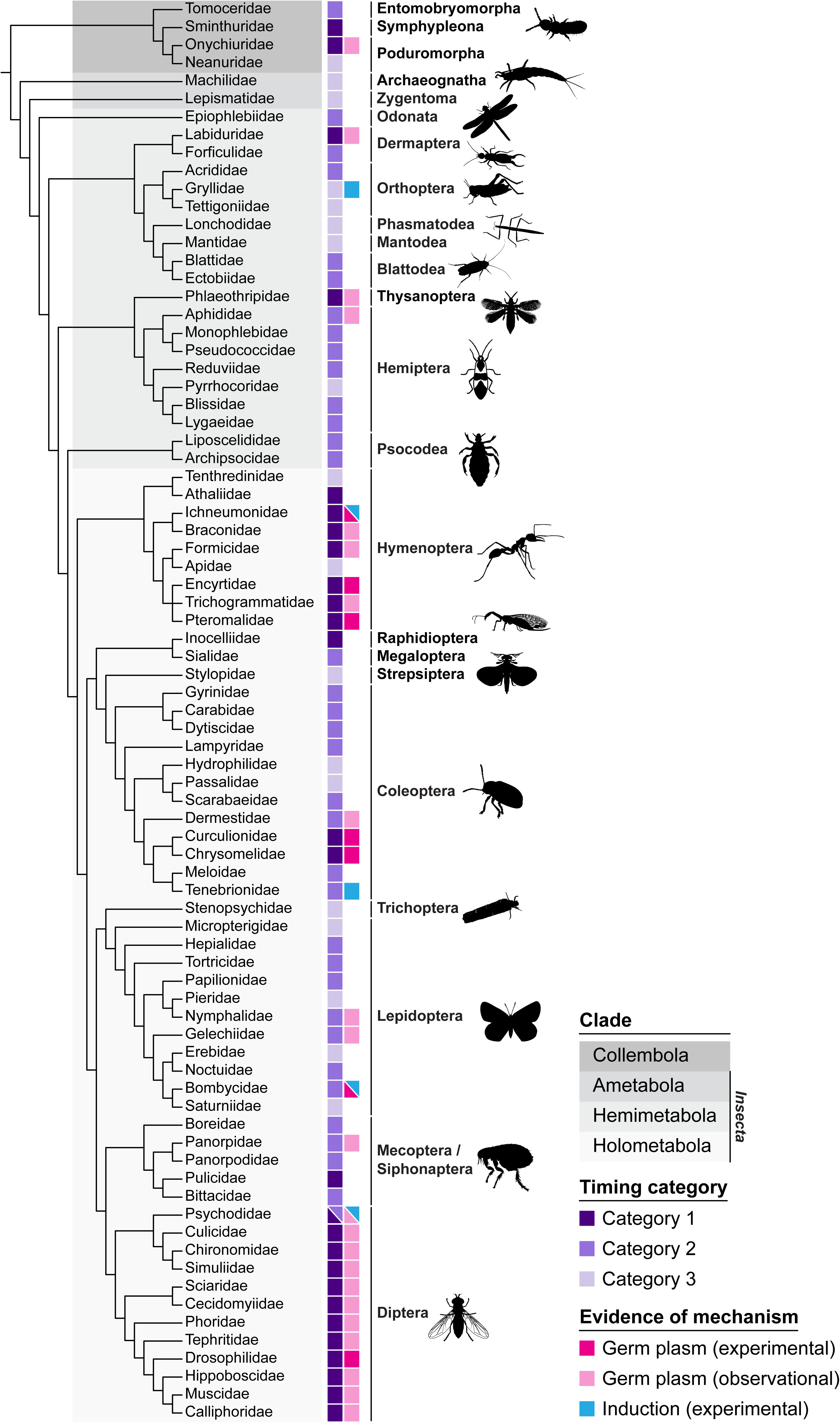
Germ cell specification across Hexapoda. Phylogeny of studied hexapod lineages indicating the PGC origin timing category and the inferred mechanism of PGC specification. Phylogenetic relationships in the cladogram are derived from published relationships listed in Table S2; branch lengths are not to scale. Data on the timing of germ cell specification are shown in the first column as follows: dark purple, Category 1 (cleavage/blastoderm); medium purple, Category 2 (gastrulation); light purple, Category 3 (mesoderm differentiation). Data on the mechanism of germ cell specification, inferred as described in Supplementary Information, are shown in the second column: dark pink, experimental evidence supports the use of germ plasm; light pink, morphological evidence of germ plasm; blue, experimental evidence of induction. Data and data sources are described in more detail in Table S1.

**Figure 3.**
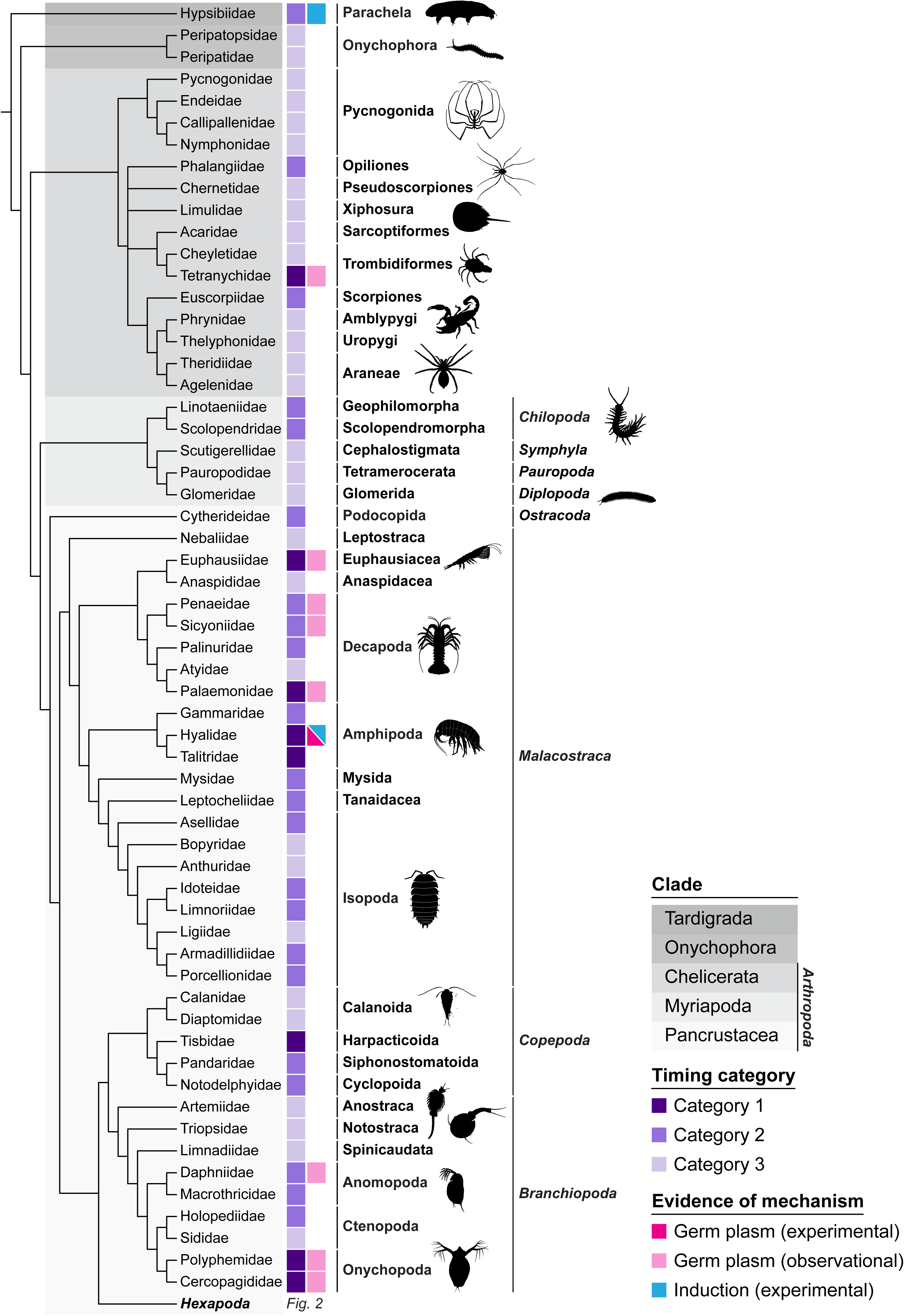
Panarthropod germ cell specification outside of Hexapoda. Phylogeny of studied non-hexapod panarthropod lineages indicating the PGC origin timing category and the inferred mechanism of PGC specification. Phylogenetic relationships in the cladogram are derived from published relationships listed in Table S2; branch lengths are not to scale. Data on the timing of germ cell specification are shown in the first column as follows: dark purple, Category 1 (cleavage/blastoderm); medium purple, Category 2 (gastrulation); light purple, Category 3 (mesoderm differentiation). Data on the mechanism of germ cell specification, inferred as described in Supplementary Information, are shown in the second column: dark pink, experimental evidence supports the use of germ plasm; light pink, morphological evidence of germ plasm; blue, experimental evidence of induction. Data and data sources are described in more detail in Table S1.

Conflicting interpretations in the literature about the timing of germ cell specification have sometimes resulted from technical advances enabling identification of germ cells at stages earlier than previously detected. For example, early work on the silk moth *Bombyx mori* (Lepidoptera) reported PGCs emerging from the mesoderm of coelomic cavities in late embryogenesis (discussed above as Category 3) (Toyama, 1902), while later studies identified PGCs on the posterior side of the ventral midline of the early germ band (as in Category 2) (Miya, 1953). The onset of PGC specification before gastrulation is further supported by data from molecular markers, which showed accumulation of *vasa* mRNA in a cluster of presumptive PGCs at the ventral midline of the early germ band (Nakao, 1999; Nakao et al., 2006). Therefore, it is possible that species which we classified here in Category 3, may be reclassified in Categories 1 or 2 by future work deploying molecular markers. This caveat notwithstanding, we note that molecular markers and functional genetic studies often corroborate late germ cell specification [see for example *Gryllus bimaculatus* (Orthoptera, Insecta) (Ewen-Campen et al., 2013b), *A. mellifera* (Hymenoptera, Insecta) (Dearden, 2006), *Parasteatoda tepidariorum* (Araneae, Chelicerata) (Schwager et al., 2015)]. We, therefore, do not assume that the hypothesis of ancestral late PGC specification is an artifact of exclusive use of morphological identification methods.

Going forward, data from key understudied lineages should be gathered to further test our hypothesis about the ancestral timing of PGC specification. For example, among the hexapods, data from members of the Palaeoptera (mayflies, dragonflies and damselflies), the ametabolous Archaeognatha and Zygentoma, or the non-insect hexapod Diplura (two-pronged bristletails) could be especially useful. Similarly, data about PGC specification are missing from Oligostraca, an understudied clade of Pancrustacea that includes Ostracoda (seed shrimp) and Branchiura (fish lice).

Outside of Pancrustacea, the field has yet to achieve clear consensus on the phylogenetic relationships of the groups within Chelicerata (Sharma and Gavish-Regev, 2025), and there is too little embryological data available for us to suggest evolutionary trends in PGC specification within this subphylum. Data regarding PGC specification are even sparser for myriapods. Specifically, we are aware of relevant data from only five species across the four myriapod classes (Dohle, 1964; Green and Akam, 2014; Heymons, 1901; Tiegs, 1940; Tiegs, 1947). Studies of PGCs in a broader range of both chelicerates and myriapods, as well as detailed embryological studies in unexplored taxa, will therefore be needed to fully understand the developmental context and evolutionary trends of germ cell specification in these subphyla.

#### Molecular and genetic mechanisms of germ cell specification in panarthropods

Beyond studying where and when germ cells first appear during arthropod embryogenesis, researchers have sought to understand how they are specified. Germ cell specification in animals has historically been classified into two mechanisms, often referred to as maternal inheritance and zygotic induction (Extavour and Akam, 2003; Nieuwkoop and Sutasurya, 1981). In maternal inheritance, germ cell determinants accumulate in a specific region of the oocyte cytoplasm, forming a germ plasm. Cells that inherit sufficient germ plasm components after the embryo divides become PGCs. In embryos that undergo zygotic induction, all cells are in principle competent to become germ cells at the beginning of embryogenesis. Certain cells later receive signals from neighboring cells, produced by the activity of the zygotic genome, which induce them to take on germ cell identity. In the following section, we describe how historical observational data and perturbational experimental approaches have offered increasingly robust evidence to distinguish between these mechanisms in several panarthropod species.

### Germ cell specification through maternal inheritance

In the early twentieth century, researchers speculated that the electron-dense, membraneless granules frequently found in the cytoplasm of PGCs were responsible for their specification. Experiments in chrysomelid beetles provided early evidence that the posterior cytoplasm of the oocyte or early embryo contains germ cell determinants. Burning or puncturing the posterior poles of beetle eggs with needles to damage or remove posterior cytoplasm reduced or eliminated PGCs (Hegner, 1908; Hegner, 1911), demonstrating that this cytoplasm does not simply correlate with PGC specification, but rather plays a causal role in PGC specification. Studies in *D. melanogaster* offered further evidence for the necessity of the germ plasm in PGC specification. Removing or destroying the posterior cytoplasm of *D. melanogaster* eggs via ultraviolet irradiation demonstrated that this region is required for germ cell formation (Geigy, 1931). Subsequent centrifugation experiments in *D. melanogaster* embryos revealed that shifting the granular structures in posterior cytoplasm away from the posterior pole prevented germ cell formation or led to germ cells forming in a different part of the embryo (Jazdowska-Zagrodzińska, 1966).

Experiments transplanting specific regions of cytoplasm between *D. melanogaster* embryos showed that the contents of the posterior cytoplasm were sufficient to specify germ cells. Transplanting posterior cytoplasm from undamaged *D. melanogaster* embryos into sterile ones in which the posterior pole had been damaged with ultraviolet irradiation rescued fertility, while transplanting anterior cytoplasm did not (Okada et al., 1974). Further transplantation experiments demonstrated that posterior pole plasm introduced into the anterior of another embryo caused formation of ectopic pole cells that gave rise to functional germ cells when transplanted again to the posterior of a third, sterile embryo (Illmensee and Mahowald, 1974). Similar experiments transferring the posterior cytoplasm from late-stage oocytes into embryos revealed that the oocyte pole plasm could specify functional PGCs (Illmensee et al., 1976), demonstrating that the necessary germ cell determinants in *D. melanogaster* are deposited during oogenesis. These foundational studies showed that maternally derived cytoplasmic granules are both necessary and sufficient for germ cell formation in some species.

Contemporary studies testing for maternal inheritance often ask whether known germ line gene products accumulate during late oogenesis or early embryogenesis in the region where PGCs first arise. For example, antibody staining or *in situ* hybridization experiments to detect germ line molecules including Vasa protein or *nanos* mRNA have suggested the presence of germ plasm in mosquitoes (Juhn and James, 2006; Juhn et al., 2008), scuttle flies (Wotton et al., 2014), ants (Khila and Abouheif, 2008; Lynch et al., 2011; Rafiqi et al., 2020), wasps (Kharel et al., 2024; Lynch et al., 2011), weevils (Lynch et al., 2011), aphids (Chang et al., 2006; Chang et al., 2007; Chen et al., 2009; Lin et al., 2014), amphipods (Gupta and Extavour, 2013) and prawns (Chen et al., 2021). Transcriptomic analyses of different portions of embryos and oocytes (Carter et al., 2015; Klomp et al., 2015; Quan and Lynch, 2016; Yoon et al., 2019) have also revealed asymmetric localization of molecules associated with PGC specification.

However, expression data showing asymmetric localization of putative germ plasm components are not sufficient to demonstrate that these molecules function in PGC specification. To support such claims, functional genetic techniques are needed to manipulate gene activity and test the effect on germ plasm assembly and germ line development. The function of germ plasm components in panarthropods has been most thoroughly studied in *D. melanogaster* (Lehmann, 2016; Mahowald, 2001; Trcek and Lehmann, 2019). The membraneless ribonucleoprotein granules at the posterior of *D. melanogaster* oocytes and early embryos are nucleated by the protein Oskar, which is both necessary and sufficient for germ plasm assembly (Ephrussi and Lehmann, 1992; Kistler et al., 2018). Oskar-mediated localization of core germ line molecules such as Vasa, Tudor, Aubergine and *nanos* to germ granules is essential for PGC specification and maintenance of germ cell identity during cell migration and gonad formation in these fruit flies (Lehmann, 2016; Mahowald, 2001).

Analysis of germ plasm composition and localization in species beyond *D. melanogaster* has revealed both conserved and rapidly evolving features. Within the *Drosophila* genus, germ granule size, morphology (Counce, 1963; Mahowald, 1968; Rivard et al., 2025), and the relative proportions of different granule components vary widely (Doyle et al., 2023). The wasp *N. vitripennis* also has a well-characterized germ plasm in the form of a single large structure called the oosome (Bull, 1982). Akin to *D. melanogaster, N. vitripennis* requires *oskar* to assemble a germ plasm containing many of the same molecules (e.g., Vasa, Tudor, and *nanos*) (Lynch and Desplan, 2010; Lynch et al., 2011). However, while *D. melanogaster* has many small germ granules (Mahowald, 1962), the *N. vitripennis* oosome has a distinct protein organization and transcript composition compared to that observed in *D. melanogaster* (Kemph et al., 2025; Kharel et al., 2024; Lynch et al., 2011). In several crustacean species, the germ plasm appears as a large, round granule in the cortex of embryos and contains conserved germ plasm markers (Chen et al., 2021; Grattan et al., 2013; Gupta and Extavour, 2013). Studying germ plasm in additional species is needed to clarify whether and how variation in germ plasm makeup and organization influences the post-transcriptional regulation and function of germ line determinants.

### Germ cell specification through zygotic induction

Experimentally demonstrating inductive germ cell specification, requiring physical, molecular or genetic perturbation methods, is more technically challenging than detecting germ plasm accumulation, for which in situ hybridization techniques or protein detection may suffice. Consequently, most studies have inferred or hypothesized zygotic induction based on absence of evidence of germ plasm. For example, *in situ* hybridization experiments fail to reveal asymmetric localization of putative germ line gene transcripts in oocytes and early embryos of various species, including milkweed bugs (Ewen-Campen et al., 2013a), crickets (Ewen-Campen et al., 2013b), honeybees (Dearden, 2006), mites (Dearden et al., 2003), spiders (Schwager et al., 2015), centipedes (Green and Akam, 2014) and tardigrades (Heikes et al., 2023). Instead, key germ line genes are expressed in a subset of cells later in embryogenesis, coincident with reports of the first PGCs based on morphological criteria (Table S1). However, we note that even in the well-accepted case of mouse PGC induction (Saitou and Yamaji, 2010), it remains formally possible that some unknown, and therefore untested, germ line determinant or factor that makes cells competent to respond to later germ line induction signals, is asymmetrically localized and preferentially inherited by germ cells.

This caveat notwithstanding, there are multiple examples of experimental support for induction rather than inheritance across panarthropods. Maternal knockdown of putative germ plasm components in milkweed bugs and crickets reveals that the maternal products of these genes are dispensable for PGC formation (Ewen-Campen et al., 2013a; Ewen-Campen et al., 2013b). Laser ablation of single cells in 2- and 4-cell stage embryos of the tardigrade *Thuliana stephaniae* does not prevent PGC specification (Hejnol and Schnabel, 2005). This observation suggests that this species lacks a cytoplasmic region with inherited germ line determinants, and that it might instead specify PGCs through inductive pathways.

Germ cell formation in embryos with tissue duplications can also be interpreted as evidence against germ plasm and consistent with induction. For example, in the drain fly *Clogmia albipunctata* and the flour beetle *T. castaneum*, genetic manipulations producing double abdomen embryos lead to germ cell formation at both anterior and posterior ends (Ansari et al., 2018; Yoon et al., 2019). Similarly, naturally occurring “twins” of the stick insect *Carausius morosus* are duplicated embryos fused along their anterior–posterior (A–P) axis, which both possess germ cells (Cavallin, 1971). In contrast, double abdomen embryos of *D. melanogaster* only produce germ cells at the true posterior end, since germ plasm is absent at the ectopic posterior end (Lasko and Ashburner, 1990).

Among panarthropods, experiments in the cricket *G. bimaculatus* have provided the most direct evidence for zygotic induction of PGCs. Knockdown of the mesoderm specification factor *twist* leads to loss of PGCs, suggesting that the mesoderm contributes to germ cell specification and/or maintenance (Ewen-Campen et al., 2013b) and consistent with morphological evidence that orthopteran PGCs first arise from the embryonic mesoderm (Wheeler, 1893). The bone morphogenetic protein (BMP) signaling pathway, which induces germ cells in mice (Hopf et al., 2011; Lawson et al., 1999; Ying and Zhao, 2001; Ying et al., 2000), is also required for cricket germ cell specification. Specifically, BMP signaling and the downstream transcription factor Blimp-1 induce a subset of mesodermal cells to become PGCs in crickets (Donoughe et al., 2014; Nakamura and Extavour, 2016).

The shared use of both BMP and Blimp-1 in mice and crickets may have convergently evolved, or could suggest that this signaling mechanism is ancestral in bilaterian inductive PGC specification (Lochab and Extavour, 2017). Determining whether BMP and its downstream target genes are required for germ line specification in more species will be necessary to distinguish between these possibilities. Only a small number of signaling pathways contribute to developmental fate decisions in animals (Pires-daSilva and Sommer, 2003), increasing the likelihood of evolutionary convergence and limiting the number of candidates to evaluate for roles in PGC induction in different species.

### Intermediate and flexible strategies of germ cell specification

Evidence from some animal species suggests that there may not be a strict dichotomy between inheritance and induction (discussed by Seervai and Wessel, 2013). These two mechanisms might instead represent ends of a spectrum that are not necessarily mutually exclusive. Below, we discuss cases of panarthropod species for which data suggest that their mechanisms of germ cell specification may fall somewhere along this continuum.

In the silk moth *B. mori*, historical studies have generated conflicting evidence about how germ cells are specified. Germ granules were not detected in early light microscopy experiments, but germ cells failed to form after cauterization of the ventral side of cleavage stage embryos, suggesting this region could contain germ cell determinants (Miya, 1958a). Subsequent studies found that Vasa protein is not asymmetrically localized before cellularization, again suggesting a lack of germ plasm, or at least a germ plasm lacking Vasa protein (Nakao et al., 2006). However, the mRNA of one of the four *B. mori nanos* homologs, *nanosO*, localizes along the ventral midline of syncytial embryos immediately following egg-laying, and is inherited by the PGCs (Nakao et al., 2008) suggesting the possibility that a germ plasm containing *nanosO* transcript specifies PGCs. Intriguingly, researchers found that maternal and zygotic *nanosO* act redundantly in PGC specification; loss of one or the other reduces but does not completely eliminate germ cells, whereas loss of both prevents germ cell formation (Nakao and Takasu, 2019). The possibility that both inheritance and induction mechanisms contribute to *B. mori* PGC specification suggests that this species could offer insight into transitions in germ cell specification mechanisms within Lepidoptera or across insects more broadly.

Although in *D. melanogaster*, PGC specification is traditionally characterized as driven by germ plasm, zygotic signaling is also suggested to contribute to germ cell specification in this species (Colonnetta et al., 2022). Loss-of-function mutations in the BMP family morphogen Dpp lead to partial loss of transcriptional quiescence and mislocalization of germ plasm in the newly formed pole cells (Colonnetta et al., 2022). Furthermore, loss of BMP signaling leads to activation of the normally suppressed terminal signaling pathway in pole cells, which in turn disrupts their migration (Colonnetta et al., 2022). These observations illustrate a universal conceptual challenge in the field of developmental biology: distinguishing between, or even consistently defining, cell type specification and maintenance. Although pole cells can form upon disruption of BMP signaling, they do not contain the proper molecular components to act as PGCs (Colonnetta et al., 2022). We suggest that it may be most useful to define PGC specification mechanisms as those that ensure both the physical formation of germ cell precursors (including adoption of specific morphologies and activation of characteristic gene regulatory states), as well as the competence of those cells to take the next developmental steps toward joining the gonad. Under this definition, we would indeed concede that both germ plasm and zygotic inductive mechanisms play roles in *D. melanogaster* germ cell specification.

The ability to post-embryonically regenerate a germ line initially specified via maternal inheritance has also been reported in multiple invertebrate species, including the sea squirt *Ciona intestinalis* (Takamura et al., 2002), the earthworm *Enchytraeus japonensis* (Tadokoro et al., 2006) and the marine worm *Capitella teleta* (Dannenberg and Seaver, 2018) (see also Özpolat, 2024 for a review of this phenomenon in Annelida). The wasp *Pimpla turionellae* (Hymenoptera) has an oosome (Meng, 1968) that is required for germ cell specification. Damaging the oosome results in no pole cells, which are the PGCs of this species (Achtelig and Krause, 1971). However, oosome-deficient embryos lacking pole cells develop into larvae with germ cells in their gonad, indicating that an additional mechanism must regenerate germ cells later in development (Achtelig and Krause, 1971). Evidence of germ line regeneration has also been reported in the amphipod *Parhyale hawaiensis* (Kaczmarczyk, 2014), which has a germ plasm that is asymmetrically segregated to the *g* micromere and its lineage (Extavour, 2005; Gerberding et al., 2002; Gupta and Extavour, 2013). Ablation of the *g* micromere eliminates PGCs in the embryo (Extavour, 2005), but the germ line is replaced post-embryonically with germ cells that appear to be derived from the mesoderm (Kaczmarczyk, 2014). The authors of this study speculated that these replacement germ cells specifically derive from cells of the somatic gonad (Kaczmarczyk, 2014), reminiscent of a previously popular theory that primordial germ cells in most animals derived from the inner epithelium of the somatic gonad (Everett, 1944; Hargitt, 1919; Heys, 1931). Species capable of regenerating PGCs present the opportunity to investigate the relative contributions of and potential interactions between inheritance- and induction-based mechanisms within a single species. Furthermore, these species may represent living examples of the intermediates that are hypothesized to exist in evolutionary transitions between modes of germ line specification (Extavour, 2007).

#### Evolutionary drivers of panarthropod germ cell specification

In the following section, we discuss the evolutionary dynamics of germ cell specification mechanisms across Panarthropoda, and propose potential developmental and genetic factors that might contribute to these dynamics.

### Convergent evolution of panarthropod germ plasm

The phylogenetic distribution of germ cell specification mechanisms across panarthropods reveals a complex evolutionary history. Data for tardigrades, chelicerates and myriapods are consistent with induction as the ancestral mechanism of panarthropod primordial germ cell specification (Figure 3), and this hypothesis is strongly supported by statistical tests described below (Figs. S1-S2). In Pancrustacea, the most heavily represented group in our dataset (Table S1), the late origin of primordial germ cells in several species suggests induction may be widely used. However, there appear to be independent origins of germ plasm in Malacostraca and Branchiopoda, a hypothesis that is also supported by formal statistical testing described below (Figs. S1-S2; Table S3). The use of induction and germ plasm in different decapod species suggests that further sampling could reveal patterns in the evolution of germ cell formation within this order (Biffis et al., 2009; Chen et al., 2021; Terao, 1929). However, functional experiments and broader taxonomic sampling will be necessary to confirm mechanisms and reconstruct their evolutionary trajectories across Panarthropoda as a whole.

Reminiscent of a broader pattern that we and others previously reported across animals (Extavour, 2007; Extavour and Akam, 2003; Johnson et al., 2003; Whittle and Extavour, 2017), in some insect orders we noted that lineages branching closer to the base of a clade deployed induction, while later branching lineages appeared to use germ plasm (Figs. 2–3). This pattern is well exemplified in Coleoptera, (Kobayashi et al., 2006; Komatsu and Kobayashi, 2012; Krause, 1947; Niikura et al., 2017; Schröder, 2006; Ullmann, 1964), with pole cells forming only in Chrysomeloidea (including long-horned and leaf beetles) and Curculionidae (Brauer, 1925; Hegner, 1908; Lynch et al., 2011). These observations raise the possibility that germ plasm may not be ancestral to Coleoptera. When considered alongside the absence of germ plasm in most Lepidoptera, the reports of germ cells that first arise in the mesoderm (Category 3) in Trichoptera (Miyakawa, 1974) and of germ cells specified at gastrulation (Category 2) in Mecoptera (Suzuki, 1990), and the distinct morphologies of *N. vitripennis* and *D. melanogaster* germ plasm (see *Molecular and genetic mechanisms of germ cell specification in panarthropods* above), are consistent with a novel hypothesis: germ plasm evolved independently multiple times across insects, rather than arising only once in the Holometabola as we and others have previously proposed (Ewen-Campen et al., 2012; Ewen-Campen et al., 2013b; Lynch et al., 2011). Our previous work suggested the independent evolution of germ plasm at the phylum level (Extavour and Akam, 2003), but here, we hypothesize widespread convergent evolution of maternal inheritance-based germ cell specification within a phylum.

To test this hypothesis, we performed ancestral character reconstruction using both maximum parsimony and maximum likelihood approaches (details in Supplementary Information, Tables S2-S7 Figs. S1-S2). Both a parsimony analysis (Fig. S1) and marginal ancestral reconstructions of a maximum likelihood approach (Fig. S2; see Table S7 for marginal probabilities) provided strong statistical support for induction as the ancestral specification mechanism in Panarthropoda, Arthropoda and Insecta, with multiple independent instances of germ plasm evolution and reversion to induction (see Table S3 for a comparison of the two approaches).

We further find strong evidence that induction was the ancestral specification mechanism in Holometabola (probability 80.6%), followed by independent transitions to germ plasm in the common ancestors of Diptera and Mecoptera (or independently within Mecoptera, Siphonaptera (fleas) and Diptera); derived lineages of Lepidoptera, in the common ancestor of Chrysomelidae and Curculionidae (Coleoptera); and in the common ancestor of Hymenoptera. Outside of Holometabola, we infer independent transitions to germ plasm in Thysanoptera, Hemiptera, Dermaptera, Collembola (which may have undergone multiple independent transitions within the class), Decapoda, the last common ancestor of Hyalidae and Talitridae (Amphipoda), and in the chelicerate *Tetranychus urticae*. Collectively, these qualitative and quantitative analyses suggest a previously unappreciated dynamism in the evolutionary history of germ cell specification within Panarthropoda.

### The relationship between germ cells and somatic germ layers

In arthropods with late formation of PGCs (Category 3), these cells appear to differentiate directly from the mesoderm (Evans, 1901; Hickman, 1936; Nelson, 1915). Consistent with this hypothesis, abolishing mesoderm prevents germ cell specification in the cricket *G. bimaculatus* (Ewen-Campen et al., 2013b). Germ cell differentiation occurs concurrently with the differentiation of many other mesodermal cell types, including the somatic gonad, musculature, heart and fat body (Johannsen and Butt, 1941). We suggest that the types of mechanisms that specify germ cells are not distinct in principle from those specifying any other mesodermal cell type in these animals.

The germ layer origin of germ cells is less clear in arthropods whose germ cells appear before or very close to the time of gastrulation (Category 2). For example, PGCs in many crustaceans are described as being the first to enter the blastopore during gastrulation, and are usually considered by researchers to be derived from the mesendoderm (e.g. Cannon, 1921; Goodrich, 1939; Manton, 1928). In contrast, germ cells in Category 2 insects arise before gastrulation (Mellanby, 1935; Ullmann, 1964), consistent with the hypothesis that germ cells are distinct from the mesoderm and may not derive from any germ layer.

Crustaceans with putative germ plasm typically have stereotypical embryonic cleavage patterns and embryonic cell lineages show that the germ cell lineage is sister to the mesoderm or the mesendoderm [e.g. Amphipoda (Extavour, 2005; Gerberding et al., 2002; Scholtz and Wolff, 2002; Wolff and Scholtz, 2002), Euphausiacea (krill) (Alwes and Scholtz, 2014), Decapoda (Hertzler, 2005)], but never to the ectoderm. In addition, removing the putative germ plasm from embryos of the amphipod *P. hawaiensis* makes the *g* micromere morphologically indistinguishable from the other micromeres, which give rise to mesoderm or endoderm in unperturbed embryos (Gupta and Extavour, 2013). This result suggests that in the absence of germ cell determinants, the *g* micromere, normally the germ line progenitor, could instead adopt mesendodermal fate. This hypothesis has not been directly tested in this amphipod because gastrulation fails after ablation of the putative germ plasm (Gupta and Extavour, 2013). Together, these data suggest a close association between the germ cell and mesendodermal lineages in these crustaceans. This relationship is apparent beyond Pancrustacea, since germ cells induced late in embryogenesis arise from the mesoderm (Category 3). We thus speculate that germ cells in panarthropods may be ancestrally derived from the mesodermal lineage, consistent with our inference that inductive germ cell specification is ancestral to panarthropods (also discussed earlier).

### Evolutionary shifts in the spatial and temporal origin of PGCs

We have seen significant variation in the spatial and temporal origin of PGCs across panarthropods (Figs. 2–3; Table S1). However, regardless of when PGCs first appear, we find a common stage across many panarthropods in which PGCs are situated in bilateral segmental clusters across multiple trunk segments. Soon after this stage, the segmental clusters on each side fuse together into bilaterally symmetrical genital ridges, corresponding to the primordial embryonic gonad (Miya, 1958b; Nelsen, 1934; Seidel, 1924; Ullmann, 1964).

Most arthropods undergo “short germ” development, wherein the blastoderm tissue generates only anterior segments, with posterior segments being added sequentially to the posterior end of the germ band later in development (Krause, 1939). In contrast, the “long germ” mode of development seen in *D. melanogaster*, wherein the blastoderm tissue immediately generates all body segments, is an evolutionary novelty found only in some insects (Clark et al., 2019; Krause, 1939). In Category 1 or 2 arthropods with short germ development, germ cells are often specified before the formation of the segments that include the future embryonic gonad. In Category 2 insects, germ cells often associate with the posterior end of the germ band and then migrate toward the anterior as the germ band elongates (Goss, 1953; Hegner, 1909b; Heming and Huebner, 1994; Paz, 1958; Suzuki, 1990; Ullmann, 1964). This migratory path is so common among insects that Johanssen and Butt included it in their description of embryogenesis in a stereotypical insect (Johannsen and Butt, 1941).

The association of germ cells with the posterior end of the embryo may have had other evolutionary consequences in insects. Many insects, both long and short germ, specify the body plan early in development via maternal deposition of axial patterning determinants, independent of the mechanism of germ cell specification (reviewed in Lynch, 2019; Rosenberg et al., 2009). For example, the short germ grasshopper *Schistocerca americana* exhibits asymmetric localization of the posterior patterning determinant *nanos* in the absence of a germ plasm (Lall et al., 2003). However, insects with germ plasm that includes axial patterning determinants, such as *D. melanogaster*, often exhibit long germ development. Based on these observations, we employed phylogenetic logistic regression and Pagel’s test of evolutionary rate covariation (Pagel, 1994) to test the hypothesis that the acquisition of long germ band development influences germ cell mechanism transitions or vice versa, and found statistically significant evidence of this association using both methods (Figs. S3-S4; Table S8). Overall, this suggests that long germ band development may drive the evolution of germ plasm, which in turn may also drive transitions to long germ band development.

We further hypothesize that a spatial shift in germ cell specification to the posterior end of the embryo in Category 2 insects created a condition in which germ line determinants are localized to the same embryonic region as posterior patterning determinants (Fig. 4B). Subsequently, germ line determinants may have hitchhiked onto the asymmetric localization machinery, coupling germ cell specification to posterior patterning and thus promoting the evolution of germ plasm (Fig. 4C). Under this hypothesis, the mechanism that specifies the germ plasm of *D. melanogaster* would represent an advanced stage of this evolutionary scenario, as its germ granules include both germ line and abdominal patterning determinants (Ephrussi and Lehmann, 1992; Trcek et al., 2015; Wang and Lehmann, 1991).

1. *N. vitripennis* is an intriguing case to study the relationship between A–P patterning and germ cell specification. The posterior cytoplasm of this wasp has two populations of *nanos* mRNA: one that is incorporated into the oosome and eventually into pole cells, and another that functions in axial patterning (Lynch and Desplan, 2010). The localization of *nanos* to the oocyte posterior in this wasp depends on actin-dependent anchoring (Olesnicky and Desplan, 2007), as seen in *D. melanogaster* (Forrest and Gavis, 2003), suggesting a conserved localization mechanism between these two species. We speculate that the PGC specification mechanism used in *N. vitripennis* might be an evolutionary intermediate between the mechanism used in species where A–P patterning is entirely independent of germ cell formation (Fig. 4A), and that used in species where these two processes are fully linked (Fig. 4C). Our evolutionary hypothesis could be tested further by examining the mechanisms of localization of posterior patterning determinants and germ line determinants in more insect species to understand how existing cytoskeletal machinery for one developmental process may have been coopted for use in another.

**Figure 4.**
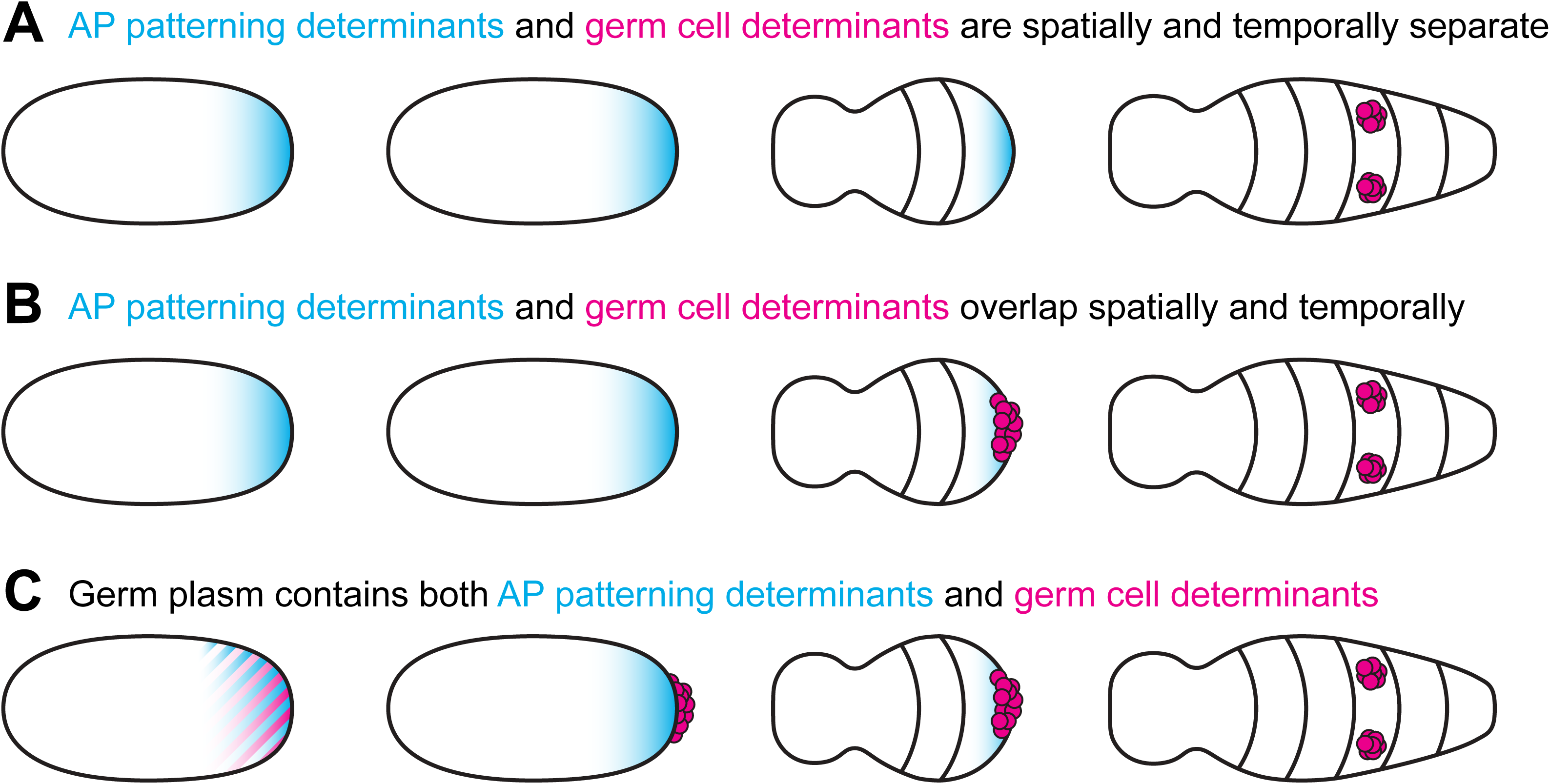
Hypothesized links between spatial and temporal dynamics of PGC origination and the mechanism of germ cell specification. (**A**) In species that induce PGCs later in embryogenesis (Category 3), the asymmetrically localized anterior–posterior (A–P) patterning determinants (blue) and germ cell determination factors (pink) are spatially and temporally separate. (**B**) Proposed shift to earlier germ cell specification at the embryonic posterior in Category 2 insects enabled overlap between the patterning and PGC determinants. (**C**) Subsequent hitchhiking of the asymmetric localization machinery by germ line determinants (indicated by mixture of blue and pink at the embryo posterior) could have coupled PGC specification and posterior patterning and thus promoted the evolution of germ plasm. Anterior is left and ventral is facing the reader in all schematics.

### Potential genetic basis of evolutionary shifts in the molecular mechanism of PGC specification

Beyond the timing and position of germ cell origin, evolutionary changes in the localization of gene products and timing of gene expression may also underlie transitions in the mechanism of specification (Extavour, 2007; Srouji and Extavour, 2010). Many of the genes important for specifying germ cells are conserved across metazoans (Ewen-Campen et al., 2010), but they exhibit distinct temporal and spatial expression patterns across species. In species that use inheritance, the germ line gene products are expressed during oogenesis and persist in the early embryonic cytoplasm to be inherited by the PGCs. In species that induce germ cells, these genes are either not expressed in oocytes or the gene products are neither asymmetrically nor preferentially inherited by PGCs. As we have suggested previously (Extavour, 2007; Srouji and Extavour, 2010), relatively simple regulatory modifications affecting post-transcriptional regulation, the cytoskeleton, or RNA or protein localization could cause germ line gene product retention in oocytes, thereby enabling the transition from induction to inheritance.

The insect-specific gene *oskar* offers a potential example of gene network evolution driving shifts in germ cell specification mechanism. Since *oskar* is likely a nucleator of germ plasm in multiple insects (Ephrussi et al., 1991; Jones and Macdonald, 2007; Kemph and Lynch, 2022; Lynch et al., 2011; Rivard et al., 2025) and has been lost multiple times since its origin before the divergence of Pterogyta from Ametabola (Blondel et al., 2021), we hypothesized that *oskar* gene presence in the genome might be correlated with germ line specification mode in a given taxon (Fig. S3). A related hypothesis, that the evolutionary advent of the *oskar* gene facilitated the evolution of holometabolous germ plasm, was previously proposed based on the co-occurrence of *oskar* transcript and germ plasm in *N. vitripennis* and *D. melanogaster*, as well as the absence of both *oskar* homologs and germ plasm in *T. castaneum*, *A. mellifera*, *C. albipunctata*, most Lepidoptera, and the hemimetabolous genomic resources available at the time (Lynch et al., 2011). The results of applying phylogenetic logistic regression to this hypothesis suggest a strong and statistically significant association between *oskar* presence and maternal inheritance (Fig. S3; Table S9). We emphasize that this observed association does not imply that *oskar* is necessary or sufficient for germ plasm formation; indeed, there are insect species without *oskar* that nevertheless have germ plasm and species with *oskar* that lack germ plasm (Fig. S3), as we discuss below.

In contrast to prior hypotheses from our group and others of a single origin of germ plasm at the base of the Holometabola (Ewen-Campen et al., 2012; Ewen-Campen et al., 2013b; Kemph and Lynch, 2022; Lynch et al., 2011), our current likelihood and parsimony reconstruction of induction as ancestral to Holometabola (Fig. 5; Figs. S1–S2) implies that *oskar* has been independently recruited as a germ plasm nucleator at least twice, in Hymenoptera (Lynch et al., 2011) and in Diptera (Ephrussi and Lehmann, 1992). The strong association between *oskar* presence and germ plasm (Fig. S3) implies that it may have been co-opted for germ plasm nucleation even more times across insects. We note that such “gene reuse” is a common feature of convergent evolution across multiple biological processes (Chan et al., 2010; Martin and Orgogozo, 2013; Morris, 2009; Reed et al., 2011; Satterlee et al., 2024). Since *oskar* is co-expressed with germ plasm components in nervous systems of hemimetabolous and holometabolous insects (Ewen-Campen et al., 2012; Kulkarni et al., 2023; Xu et al., 2013), we posit that independent acquisitions of germ plasm in distinct lineages may have proceeded via parallel cooption of a pre-existing gene interaction network. Future studies should aim to determine whether each independent origin of germ plasm in the Holometabola arose by co-option of *oskar* to the germ line, or by *oskar*-independent mechanisms of germ plasm assembly.

**Figure 5.**
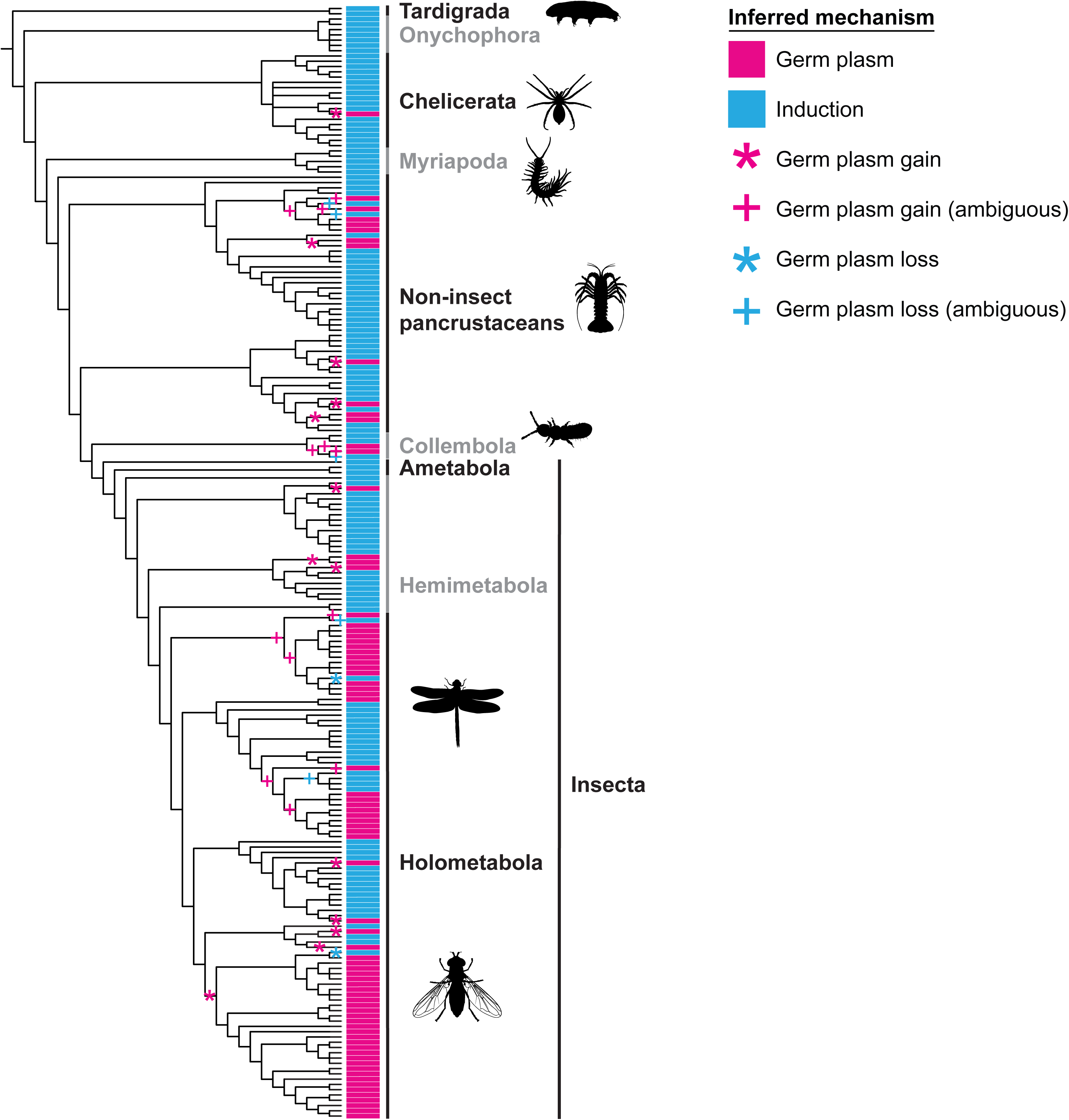
Evolutionary dynamics of panarthropod germ cell specification. Phylogenetic relationships among all study species (Table S1) are derived from published relationships listed in Table S2; branch lengths are not to scale. Additional taxonomic information is provided in Fig. S1. Colored lines to the right of each branch in the cladogram designate the mechanism of germ cell specification (inferred as described in Supplementary Materials) as follows: pink, germ plasm; blue, induction. Pink asterisks (*) denote proposed instances of germ plasm evolution (based on maximum parsimony) and pink pluses (+) mark ambiguous cases of germ plasm gain. Blue asterisks (*) denote proposed instances of germ plasm loss and blue pluses (+) mark ambiguous cases of germ plasm loss.

The results of our analysis (Fig. S3; Table S9) suggest that *oskar*’s functional evolution may have contributed to a transition from induction to inheritance, but precisely when and how *oskar* acquired a germ line role remains unclear. In the genomes of species of Carabidae (ground beetles), a beetle family that lacks pole cells, *oskar* orthologs have been identified, decoupling the presence of *oskar* from germ plasm in Holometabola (Blondel et al., 2021; Kobayashi et al., 2013). Morphological data suggest that thrips use germ plasm (Haga, 1985; Heming, 1979), and *oskar* homologs have been found in this order (Blondel et al., 2021). If *oskar* plays a role in thrips germ cell specification, it could support the entirely novel hypothesis that *oskar* evolved its germ line determinant properties before the evolution of complete insect metamorphosis.

### The case for novel panarthropod germ plasm nucleators

Recruitment of *oskar* to the germ line is not the only mechanism explaining the evolution of germ plasm in arthropods. The genome of the hemimetabolous pea aphid *Acyrthosiphon pisum* lacks an *oskar* ortholog but their embryos have a germ plasm containing Vasa and Nanos proteins (Chang et al., 2006; Chen et al., 2009; Lin et al., 2014), which are asymmetrically localized by unknown mechanisms. Vasa is also asymmetrically localized to a putative germ plasm in amphipod and decapod species (Chen et al., 2021; Gupta and Extavour, 2013), which is likely to have evolved via distinct mechanisms since *oskar* is an insect-specific gene (Blondel et al., 2021). Outside of arthropods, germ plasm nucleators have convergently evolved in diverse animal phyla (Aoki et al., 2016; Bontems et al., 2009; Kulkarni and Extavour, 2017; Lee et al., 2020; Marlow and Mullins, 2008; Scholl et al., 2024). Similarly, we posit that germ plasm nucleators other than *oskar* likely also exist across the panarthropods, awaiting discovery.

## Concluding remarks

By reinterpreting the meticulous work of generations of classical embryologists through the lens of modern developmental genetics and an improved panarthropod phylogeny, we have generated four major, novel and testable evolutionary hypotheses about the mechanisms that specify the germ line. First, inductive mechanisms likely specified PGCs ancestrally in Panarthropoda, and differentiation from the mesoderm is the likely ancestral mode of PGC specification in Euarthropoda. Second, germ plasm arose independently multiple times across Panarthropoda, in many different crustacean and insect orders. Third, in contrast to the *Drosophila* paradigm of PGC specification via germ plasm, induction was likely the ancestral PGC specification mechanism in the Holometabola, and germ plasm likely arose independently in some lineages of flies, fleas, beetles, wasps, ants, and moths. Finally, the *oskar* gene was likely co-opted as a germ plasm nucleator at least twice independently in flies and wasps, and many panarthropods are likely to use something other than *oskar* to nucleate their germ plasm. Together, these hypotheses have implications for the genetic basis of convergence in evolution that extend beyond the specific problem of germ cell formation. The advent of new genetic tools in an increasing number of panarthropod model species places us in an exciting era to test the new hypotheses we propose here, tackle age-old questions about the evolution of panarthropod germ cell specification, and understand the impact of these mechanisms of genome evolution and body patterning in the most successful group of animals on earth.

## Acknowledgements

We thank members of the Extavour Lab for helpful discussions, Kira Heikes for the image of tardigrade germ cells (Fig. 1B4), and Prashant Sharma for the image of harvestmen germ cells (Fig. 1B3).

## Data Availability

All data sources are listed in the accompanying Supplementary Material and/or are available at https://github.com/rishabhrajkapoor/panarthropoda_gc_specification_evolution, commit ID ff9945c.

## Author Contributions

All authors gathered data and performed data analysis and interpretation. JAK, ELR and RRK wrote the first draft of the manuscript. CGE conceived of the study, obtained funding for the study, supervised its execution, and reviewed and edited the manuscript.

