## Supplementary Information for "The dynamic evolution of panarthropod germ cell specification mechanisms"

**There and back again: the dynamic evolution of panarthropod germ cell specification mechanisms**

**This document contains the following materials:**

- Supplementary Methods
- Supplementary Figures S1 through S5
- Supplementary Tables S1 through S9
- Supplementary References

**Supplementary Methods**

Throughout these Methods, custom scripts (blue text) and output files and folders (magenta text) described are located at the github repository for this study, <https://github.com/rishabhrajkapoor/panarthropoda_gc_specification_evolution>, commit ID ff9945c. Analysis packages used are indicated in green text. Publicly available scripts used are indicated in gold text.

*Inferring germ cell specification mechanisms from published literature*

For several species, experimental manipulations have provided direct evidence for the use of either germ plasm or induction (Table S1). For other species with only observational data available in published literature, we inferred the likely mechanism of specification for the ancestral character reconstruction based on patterns in the developmental timing and location of germ cell origination. For species designated as Category 1, with germ cells forming in the early blastoderm and cleavage stages, we inferred that they were using germ plasm. For species designated as Category 3, with germ cells forming late in embryogenesis from the mesoderm, we inferred that they were using induction. For Category 2 species, we inferred the use of germ plasm for species in which granules with the characteristic morphological and spatiotemporal appearance of germ granules (e.g. Eddy, 1975) were reported; otherwise, we inferred induction. Our inferred mechanisms are available in Table S1 and in all_species_annotated.tsv.

### *Phylogram construction*

To visualize the phylogenetic relationships among the 211 species in our dataset of germ cell specification data (Table S1), we concatenated published phylogenies from 23 pre-existing studies of various panarthropod lineages (Table S2) (using notebook phylogram_constraints.ipynb, output file constraint_trees/all_combined_species.nwk). Newick trees were obtained directly from the published studies, from the TimeTree (Kumar et al., 2022) or from the Open Tree of Life (OpenTreeOfLife et al., 2019) websites when available, or were manually extracted from published figures. Individual Newick trees are available in the “constraint_trees” directory. Phylogenetic relationships were delimited up to the level of taxonomic family, except for the families Chrysomelidae, Culicidae, and Drosophilidae, which were further resolved to the subfamily, genus, and species levels, respectively, due to the greater representation of these families in our dataset. We used polytomies to represent intrafamily relationships, paraphyly in inter-family relationships, or unresolved nodes in the arthropod phylogeny. Phylograms were taxonomically annotated with the ete3 package (Huerta-Cepas et al., 2016), manually concatenated, and visualized in the iTOL web server (Letunic and Bork, 2024). Branch lengths in the resulting phylograms (Figures 2-3, Figure S1) are arbitrary and do not reflect evolutionary divergence times.

### *Ancestral character reconstruction via maximum parsimony*

Ancestral character reconstruction of inferred germ cell specification modes (GP: germ plasm; IND: induction) via parsimony was performed on the above phylogram with the GUI interface of Mesquite version 4.0 using default parameters (Figure S1; see Table S3 for hypothesized transitions; input nexus file with GP/IND annotations available at GC_mech_parsimony_input.nex) (Maddison and Maddison, 2025). The results inferred that the ancestral panarthropod specification mode was induction, with a minimum of 24 changes over the history of the panarthropods. When considering all equally parsimonious character histories, changes from induction to maternal inheritance (min = 18, max = 23, average = 20.5) were the most common, but reverse transitions were also inferred (min = 2, max = 7, average = 4.5). At a minimum, reversions from maternal inheritance to induction were inferred in the honeybee *A. mellifera* (Hymenoptera) and the drain fly *C. albipunctata* (Diptera). We note that all maximum parsimony reconstructions assign induction as the ancestral state of Holometabola (Figure S1, node 11), which requires many fewer total transitions within Holometabola (N = 12) than would be required if germ plasm evolved a single time in the last common ancestor of Holometabola (N = 20) as previously proposed (Ewen-Campen et al., 2013a; Lynch et al., 2011) .

### *Maximum likelihood phylogenetic tree inference*

Ancestral character reconstruction requires as an input a phylogenetic tree in which branch lengths are proportional to evolutionary divergence (executed with time_calibrated_busco_phylogenomics.ipynb). To obtain genomic data for phylogenetic inference, we searched NCBI for genomes or transcriptome shotgun assemblies (TSAs) corresponding to our taxa of interest, filtering by >50% BUSCO (Manni et al., 2021) single copy complete ortholog completeness (with arthropoda_odb12, using the “Download and prep protein datasets” portion of time_calibrated_busco_phylogenomics.ipynb) (Table S4). We selected sequence data from exact species matches when available (N = 77 species), else substituted sequence data from a different species in the same taxonomic family (N = 45 substitutions) or order (N = 2 substitutions) when the germ cell specification mechanism was the same for all species in the family or order respectively in our dataset (Table S1). In the case of a substitution, the leaf name in the phylogenetic tree was replaced by the name of the family or order shared with the species in our dataset (Figure S2).

We used BUSCO version 5.8.3 with the arthropoda_odb12 set (Manni et al., 2021) to extract single-copy protein ortholog sequences. Upstream of BUSCO, protein predictions were generated for TSAs using TransDecoder version 5.7.1 (using transdecoder.sh and run_busco_prot_tsa.sh) (Haas et al., 2013) or extracted from .fa files for annotated genomes in NCBI, selecting the longest splice isoform per gene (using the “Download and prep protein datasets” portion of time_calibrated_busco_phylogenomics.ipynb and run_busco_annotated_genome.sh). For unannotated genes, protein sequences were extracted directly from nucleotide fasta files using BUSCO with the “-m genome” flag. BUSCO protein sequences were submitted to the BUSCO_phylogenomics pipeline (release 20240919) (McGowan, 2024) with default parameters to generate a concatenated supermatrix of trimmed MUSCLE alignments (Edgar, 2004) (output supermatrix: BUSCO_py/supermatrix/SUPERMATRIX.phylip).

Deep-branching panarthropod relationships have been comprehensively examined by several prior studies and are often susceptible to reconstruction artifacts (Giribet, 2018; Meusemann et al., 2020). We therefore used the same literature-derived tree topologies (Table S2) used for phylogram construction to constrain the family, order, class, and phylum-level relationships in our trees when these ranks had previously been shown to be monophyletic (constraint tree at BUSCO_py/constraint_tree_iqtree_species.tree). Polytomies in our constraint tree were resolved via maximum likelihood inference from our supermatrix with IQ-Tree version 2.4 (model: LG+G4, with per-gene partitions) (Minh et al., 2020) (executed with run_iqtree_constrained_search.sh, outputs in BUSCO_py/supermatrix with the “constrained_optimized” prefix). The resulting maximum likelihood tree was rooted with Tardigrada as an outgroup (Howard et al., 2022).

### *Time calibration*

Approximate likelihood molecular clock calibrations of our IQ-Tree maximum likelihood tree were performed using the IQ2MC workflow (Demotte et al., 2025), which integrates IQ-Tree version 3.0.1 (Wong et al., 2025) with MCMCTree in PAML version 4.10.9 (Yang, 1997; Yang, 2007) (using time_calibrated_busco_phylogenomics.ipynb). The fossil calibrations were derived from references listed in Table S5, including (Wiegmann et al., 2011) for Diptera, (Misof et al., 2014) for non-dipteran insects, (Bernot et al., 2023) for non-insect Pancrustacea, and (Howard et al., 2022) for Chelicerata and the internal nodes of the panarthropod phylogeny (rooted constraint tree available at BUSCO_py/supermatrix/fossil_rooted_constrained_optimized.treefile). Hessian matrices and gradient vectors were obtained using IQ-Tree with the “-mcmctree” flag, the pre-computed supermatrix and maximum-likelihood tree, an LG+G4 substitution model, and a single partition to promote computational efficiency (using iqtree3_iq2mcmc.sh, outputs available at BUSCO_py/supermatrix/ with prefix “mcmc_prep_no_partition”). MCMCTree was then run with an independent rates relaxed molecular clock, burnin = 20000, sampfreq = 100 and nsample = 20000. Convergence of inferred divergence times was confirmed via two independent runs of MCMCTree (PAML control files at BUSCO_py/supermatrix/run1R_mcmc_prep_no_partition.mcmctree.ctl and BUSCO_py/supermatrix/run2R_mcmc_prep_no_partition.mcmctree.ctl, output files in the same directory with mcmctree.out suffix, selected time-calibrated Newick tree at mcmctree.tree).

### *Maximum likelihood ancestral character reconstruction*

We used the fossil-calibrated phylogeny of 124 species to infer ancestral states via marginal maximum likelihood using the phytools package (Revell, 2024) (using maximum_likelihood_ancestral_reconstruction.R). Five *Mk* substitution models were fit to the data, including models allowing for gamma heterogeneity in rates among branches or hidden rate transitions to account for possibly divergent evolutionary regimes under the deep time scales analyzed in our data (Table S6). The best-fitting model by the Akaike information criterion was the symmetric rates model with gamma-rates heterogeneity (Table S6), suggesting that although the overall rate of transition from germ plasm to induction is equivalent to the rate of the reverse transition, the value of this single rate parameter shows variation over the phylogeny (Revell and Harmon, 2024). The equivalent rates of transition to and from germ plasm are consistent with the reversions to induction noted in the parsimony analysis above, although we note that the quantification of evolutionary rate was only possible with our time-calibrated phylogeny. The rate heterogeneity is consistent with the observation via maximum parsimony of 15/24 transitions in Insecta alone (Figure S1). Inferred transitions with support from maximum likelihood are reported in Table S3.

*Inferring germ band development type from published literature*

We classified species by germ band type based on embryological descriptions as follows: in short germ band embryos, the initial germ band comprises only head segments; in intermediate germ band embryos, the initial germ band comprises the head and thorax; and in long germ band embryos, the initial germ band comprises head, thorax and abdominal segments.

### *Testing the relationship between germ band development type and germ cell specification mode*

To test the hypothesis that long vs. short germ band development (Krause, 1939) influences the evolution of germ plasm in insects, we first performed phylogenetic logistic regression (Ives and Theodore, 2010) with phylolm (version 2.6.5 phylolm, method = logistic_MPLE), in which the independent variable was germ band type (three-level categorical predictor with “short” as the reference level and “long” and “intermediate” as two alternatives) and germ cell specification mode was the dependent variable (induction = 0, germ plasm = 1; Figure S3) (using germ_band_correlations.R, with the phylogeny in insect_only.tree and data in insect_germ_band.tsv). We found a statistically significant association between long germ band development and germ plasm (coefficient = 3.02, p-value = 2.81e−05) and no such association between intermediate germ band development and germ plasm (coefficient = −0.01, p-value = 0.99).

Since Pagel’s test (Pagel, 1994) is performed on binary variables, we recoded all “intermediate” germ band annotations as “short” to produce a binary trait (using germ_band_correlations.R). The null model assumes that the rate of evolution of germ cell development mode is independent of the transition in long vs. short germ band development. Our three alternative hypotheses assume either that germ band type and germ cell specification mode mutually influence each other’s evolution (“bidirectional”), or that germ band type influences germ cell specification mode but not vice versa (“GC dependent”), or that germ cell specification mode influences germ band type but not vice versa (“GB dependent”). Maximum likelihood rate fits for the bidirectional model are shown in Figure S4.

All Pagel covariation models assuming coupled evolutionary rates among these two characters are significantly better fits to the data than the null model of independent evolution (likelihood ratio test p-value < 1e−3; Figure S4; Table S8). We note that the bidirectional and GB-dependent models perform similarly to the GC-dependent model.

#### *Search for* oskar *orthologs*

We searched the genomes and transcriptomes listed in Table S4 using hmmsearch (Eddy, 2023) with the consensus profile HMMs for the LOTUS (LOTUS_CONSENSUS.hmm) and OSK (OSK_CONSENSUS.hmm) domains as previously defined (Blondel et al., 2021) (using oskar_search.ipynb, hmmsearch outputs are in the hmmsearch_results folder). For this *oskar* ortholog search, a chromosome-level, unannotated genome assembly (GCA_965637365.1) for *Clogmia albipunctata* that became available at the time of performing this analysis was substituted for the genome assembly for this species that is listed in Table S4 (GCA_022818195.1). Following our previous approach (Blondel et al., 2021), we defined Oskar orthologs as proteins that were hmmsearch (E-value < 0.05) hits to both the LOTUS and OSK domains. Amino acid sequences of all detected Oskar proteins are available in oskar_proteins.fa file b.

For unannotated genomes, we first obtained *ab initio* protein predictions using the AUGUSTUS webserver with default parameters (Stanke et al., 2004). Pre-trained species parameters for gene model prediction were selected from close phylogenetic relatives in the AUGUSTUS web server as follows: *Drosophila melanogaster* for *C. albipunctata* (GCA_022818195.1) and *Mayetiola destructor* (GCA_000149185.1); *Nasonia vitripennis* for *Macrocentrus cingulum* (GCA_045786645.1); *Heliconius melpomene* for *Epiphyas postvittana* (GCA_964304675.1) and *Nymphalis antiopa* (GCA_964258955.1); *Rhodnius prolixus* for *Pyrrhocoris apterus* (GCA_039877355.1); and *Tribolium castaneum* for *Tribolium confusum* (GCA_029207805.1).

Due to the unique position of *C. albipunctata* (family: Psychodidae) as the sole dipteran lacking germ plasm in our dataset, and its lacking a published annotated genome, we additionally tested the hypothesis that this species’ genome lacks *oskar* via tblastn (blast version 2.9.0) with the *Phlebotomus papatasi* (family: Psychodidae) Oskar protein (XP_055701593.1) as a query (using oskar_search.ipynb). We found no full-length tblastn hits but did find a possible hit to the OSK domain with evidence of multiple in-frame stop codons (Figure S5), consistent with the hypothesis of pseudogenization of *oskar* in *C. albipunctata*.

#### *Testing the relationship between oskar presence and germ cell specification mode*

Since gene loss in metazoans is thought to be irreversible (Elmer and Clobert, 2025) and Pagel’s test (Pagel, 1994) permits bidirectional state transitions, we opted only for phylogenetic logistic regression with phylolm (method logistic_MPLE) to test the association between germ cell specification mode (dependent variable, 1 = germ plasm, 0 = induction) and *oskar* presence (independent variable, 1 = presence, 0 = absence) within insects while accounting for phylogenetic relatedness. We analyzed all 81 insect datasets shown in Table S9, including 24 within-family substitutions when genomic or transcriptomic data were not available for a species (using oskar_fitphyloglm.R with oskar_data.tsv and insect_only.tree as inputs).

We found no association of germ-plasm mode with BUSCO completeness (coefficient = 0.23, p-value = 0.37). However, *oskar* presence (Table S9) was moderately (although not statistically significantly) associated with BUSCO completeness (mean BUSCO completeness for *oskar*-containing datasets = 83.29%, mean BUSCO completeness for *oskar*-absent datasets = 78.09%, Mann-Whitney U p-value = 0.08). We therefore additionally included the z-score normalized BUSCO score as a covariate in our model

The results showed a significant association between *oskar* presence in the genome and the use of germ plasm to specify PGCs (coefficient = 2.58, adjusted odds-ratio: = 13.28, p-value = 0.0006). The association between *oskar* presence and maternal inheritance mode remained strong and statistically significant even after removing 24 within-family substitutions made when neither genomic nor transcriptomic data were available for a species (coefficient = 2.56, adjusted odds ratio = 12.93, p-value = 0.0003).

*Tree visualization*

Trees were visualized and annotated using the iTOL web server v7 (Letunic and Bork, 2024)**,** except for Figure S1 and Figure S2, which were generated as outputs of Mesquite and phytools, respectively.

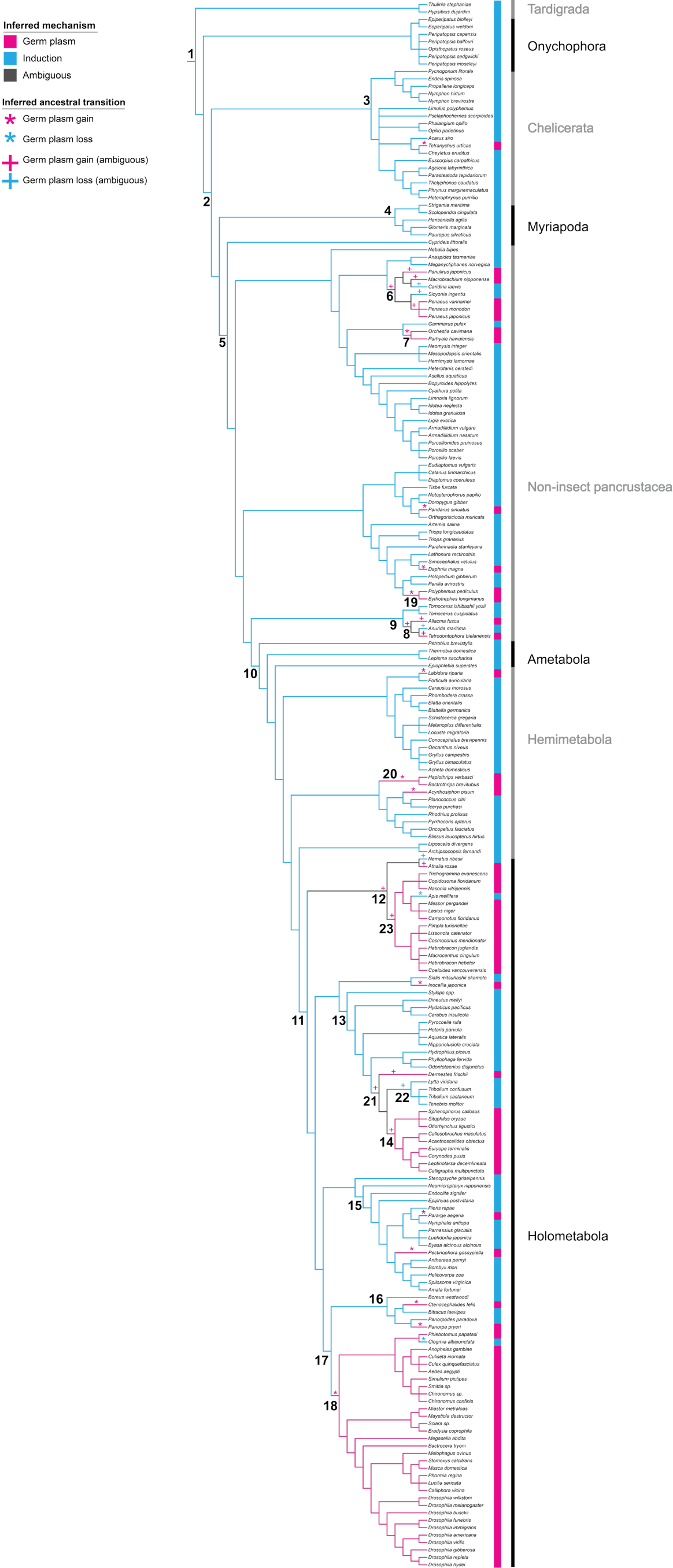

**Supplementary Figure S1 (previous page). Phylogram with maximum parsimony ancestral state inference.** Phylogram of the 211 species for which we compiled germ cell specification data in this study (Table S1), with select inner nodes numbered for reference (1 = Panarthropoda; 2 = Arthropoda; 3 = Chelicerata; 4 = Myriapoda; 5 = Pancrustacea; 6 = Decapoda; 7 = Hyalidae + Talitridae; 8 = Poduromorpha + Symphypleona; 9 = Collembola; 10 = Insecta; 11 = Holometabola; 12 = Hymenoptera; 13 = Coleoptera; 14 = Chrysomelidae + Curculionidae; 15 = Lepidoptera; 16 = Mecoptera + Siphonaptera; 17 = Antliophora (Diptera + Mecoptera + Siphonaptera); 18 = Diptera; 19 = Onychopoda; 20 = Thysanoptera; 21 = Meloidae + Tenebrionidae + Dermestidae + Chrysomelidae; 22 = Meloidae + Tenebrionidae). Species are labeled with blue or pink boxes to indicate induction or germ plasm, respectively. Internal branches are colored according to inferred ancestral states, with grey indicating that multiple equally parsimonious ancestral states are possible. Asterisks indicate unambiguous transitions (gains or losses of germ plasm); i.e., no other transition at these nodes would result in a parsimonious character history. Plus (+) signs indicate possible transitions at nodes with ambiguous character states. Note that some ambiguous character histories are mutually exclusive: e.g., one parsimonious scenario for Decapoda (node 6) is a gain of germ plasm at node 6 followed by terminal losses in the branches leading to *Caridina laevis* and *Sicyonia ingentis*, while another equally parsimonious but mutually exclusive scenario would be independent gains in *Panulirus japonicus*, *Macrobrachium nipponense,* and the common ancestor of the genus *Penaeus*.

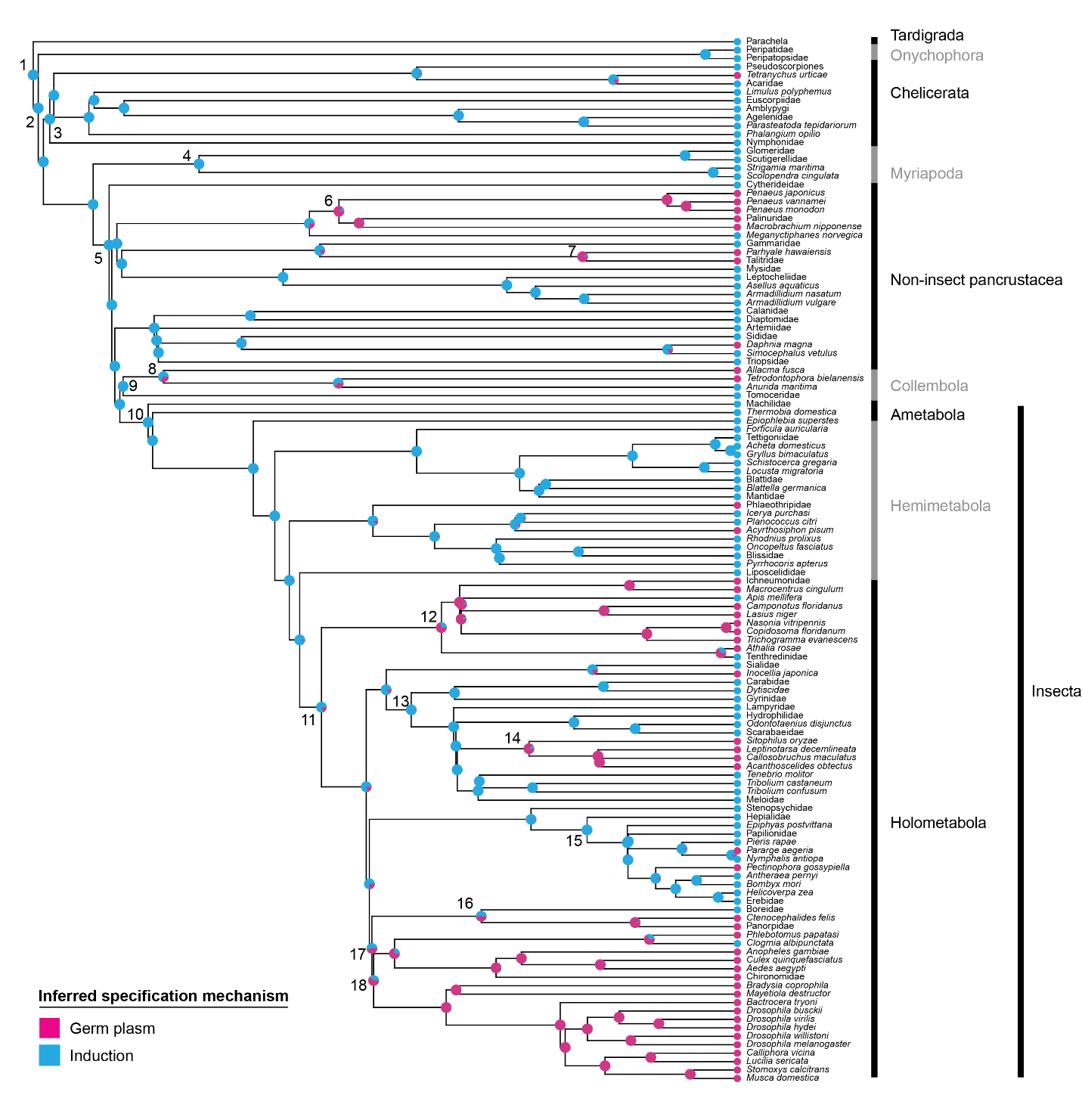

**Supplementary Figure S2. Maximum likelihood ancestral state reconstruction.** Maximum likelihood inference was performed under the symmetric transition model with gamma rate heterogeneity. Tips are labeled with colored circles according to observed induction (blue) or germ plasm (pink) mode. Pie charts at internal nodes report marginal probabilities of induction or germ plasm as the ancestral state. Select ancestral nodes are numbered with clade names and probabilities reported in Table S7. Leaf names at the family or order levels indicate a substitution for tree construction.

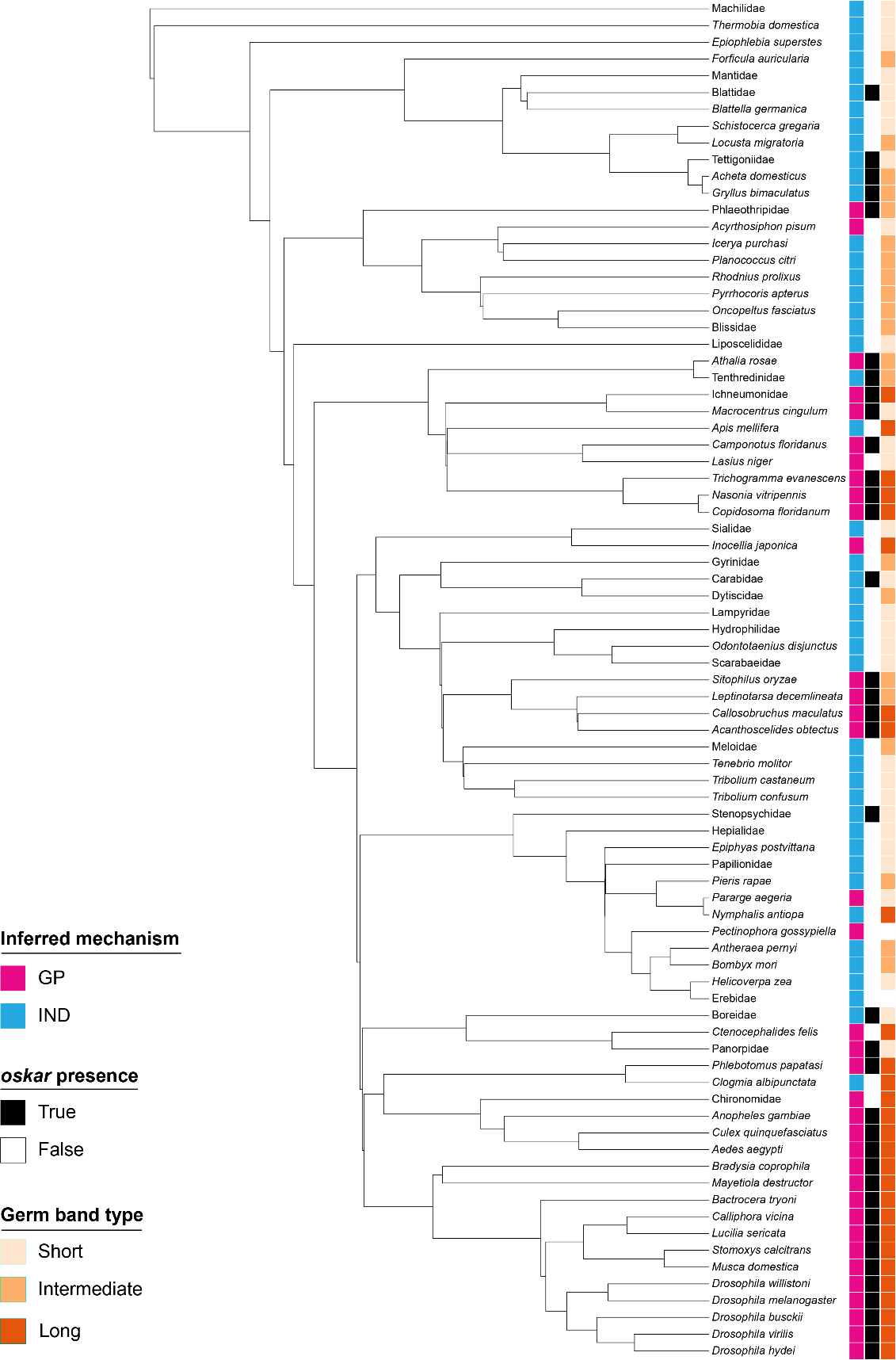

#### **Supplementary Figure S3. Relationship between germ cell specification mode (pink/blue), *oskar* presence (black/white), and germ band type (orange shades).** Time-calibrated phylogeny of the insect species with genomic/transcriptomic data available at the time of writing (N = 81; Table S9). Leaf names at the family level indicate a substitution for tree construction and inference of *oskar* presence or absence.

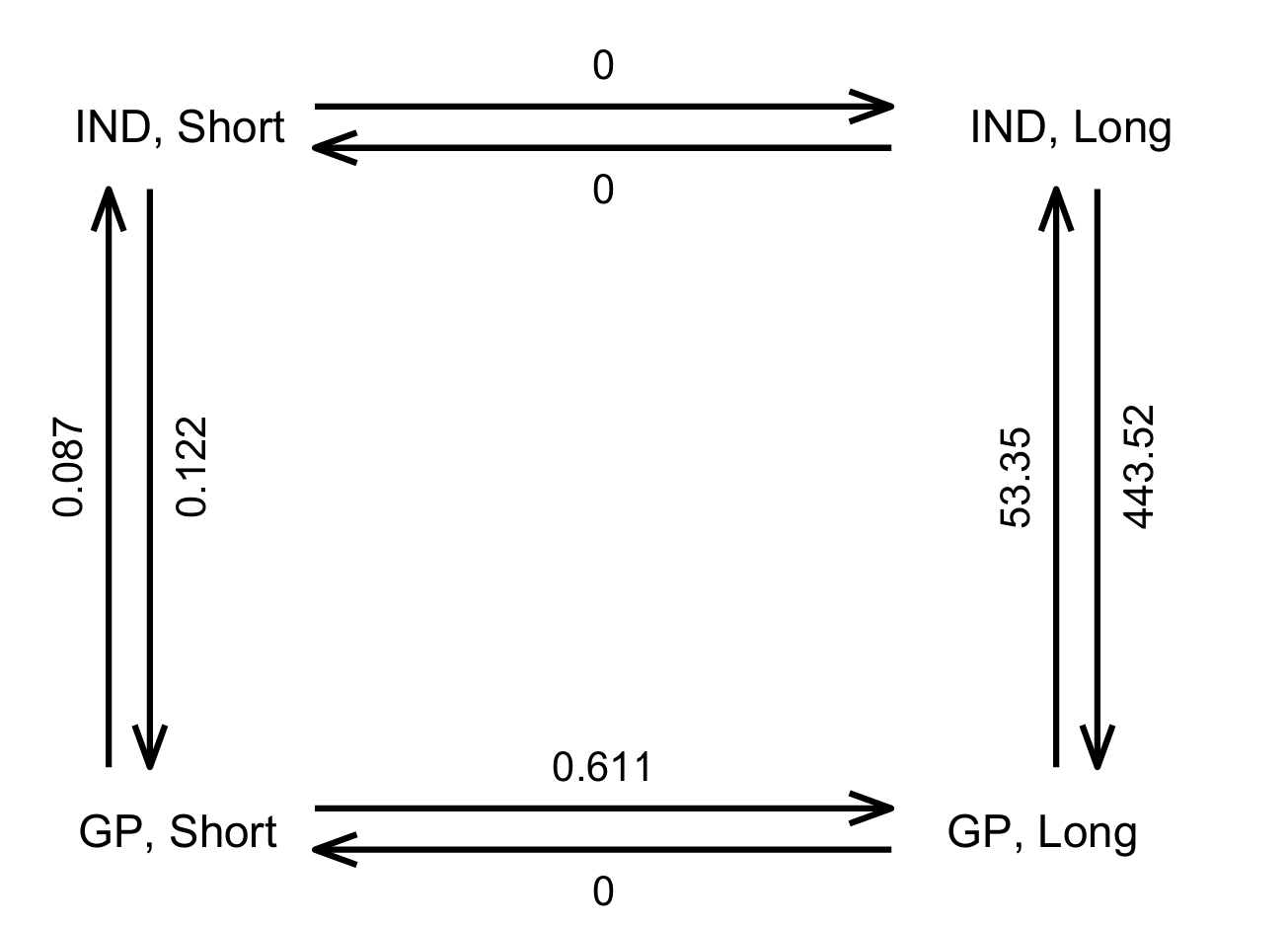

**Supplementary Figure S4. Pagel test result.** Maximum likelihood rate fits for Pagel’s test of a bidirectional dependency between transition rates in germ cell specification mode and germ band mode. GP = germ plasm; IND = induction; Short = short germ band; Long = long germ band. A relationship between germ plasm and long germ band modes is supported by the higher transition rate from “GP, Short” to “GP, Long” (0.611) than from “GP, Long” to “GP, Short” (0) or from “IND, Short” to “IND, Long” (0). Likewise, the transition rate from “IND, Long” to “GP, Long” (443.52) is higher than the transition rate from “GP, Long” to “IND, Long” (53.55). The dependent rates model better fits the data than an independent rates model (Table S8).

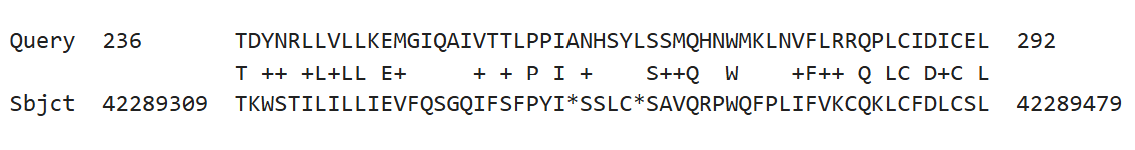

**Supplementary Figure S5. tblastn results of Oskar against the *Clogmia albipunctata* genome.** The sole tblastn hit of *Phlebotomus papatasi* Oskar protein (XP_055701593.1) against the *C. albipunctata* genome (version GCA_965637365.1), showing a partial hit on chromosome 2 (accession OZ281279.1) to the OSK domain.

**Supplementary Table S1.** Data on the timing and mechanism of germ cell origin across Panarthropoda. The “GP?” column designates whether observational (OB) or experimental (EX) data have supported the existence of germ plasm. The “IND?” column designates whether observational (OB) or experimental (EX) data have supported induction as the mechanism of germ cell specification. The “timing” column describes the timing of germ cell origination during embryogenesis and categorizes the timing as (1) blastoderm formation, (2) gastrulation, or (3) mesoderm differentiation. The “location” column describes the spatial position of germ cells when they are first reported as detectable. The “criteria” column lists the nature of the data used to identify germ cells and infer mechanism of specification: LM = light microscopy, TEM = transmission electron microscopy, SEM = scanning electron microscopy, MM = molecular markers, CL = cell lineage tracing, TS = transcriptomic data, EX = experimental manipulations.

| **order** | **suborder** | **family** | **scientific name** | **common name** | **GP?** | **IND?** | **timing** | **location** | **criteria** | **reference** |
| --- | --- | --- | --- | --- | --- | --- | --- | --- | --- | --- |
| **TARDIGRADA** | | | | | | | | | | |
| Parachela |  | Hypsibiidae | *Thulinia stephaniae* |  |  | EX | (2) cleavage | blastopore | LM, CL, EX (laser ablation) | (Hejnol and Schnabel, 2005) |
| Parachela |  | Hypsibiidae | *Hypsibius dujardini* |  |  |  | (2) cleavage | blastopore | LM, MM (*piwi*, *vasa in situ*), CL | (Gabriel et al., 2007; Heikes et al., 2023; Wenck, 1914) |
| **ONYCHOPHORA** | | | | | | | | | | |
|  |  | Peripatidae | *Eoperipatus weldoni* |  |  |  | (3) late segmented germ band | posterior germ band (walls of splanchnic mesoderm) | LM | (Evans, 1901) |
|  |  | Peripatidae | *Epiperipatus biolleyi* |  |  |  | (3) early segmented germ band | posterior germ band | TEM | (Mayer and Tait, 2009) |
|  |  | Peripatopsidae | *Opisthopatus roseus* |  |  |  | (3) early segmented germ band | posterior germ band | TEM | (Mayer and Tait, 2009) |
|  |  | Peripatopsidae | *Peripatopsis capensis* |  |  |  | (3) segmented germ band | posterior germ band | LM | (Manton, 1949; Sedgwick, 1887) |
|  |  | Peripatopsidae | *Peripatopsis balfouri* |  |  |  | (3) segmented germ band | posterior germ band | LM | (Manton, 1949) |
|  |  | Peripatopsidae | *Peripatopsis moseleyi* |  |  |  | (2) gastrulation | blastopore | LM | (Manton, 1949) |
|  |  | Peripatopsidae | *Peripatopsis sedgwicki* |  |  |  | (3) segmented germ band | posterior germ band | LM | (Manton, 1949) |
| **CHELICERATA** | | | | | | | | | | |
| Pantopoda | Colossendeoidea | Pycnogonidae | *Pycnogonum litorale* |  |  |  | (3) fifth instar | flanking the midgut | LM, TEM, SEM | (Alexeeva and Tamberg, 2021) |
| Pantopoda | Phoxychilidioidea | Endeidae | *Endeis spinosa* |  |  |  | (3) sixth instar | mesoderm surrounding midgut | LM | (Dogiel, 1913) |
| Pantopoda | Nymphonoidea | Nymphonidae | *Nymphon hirtum* |  |  |  | (3) sixth instar | mesoderm surrounding midgut | LM | (Dogiel, 1913) |
| Pantopoda | Nymphonoidea | Nymphonidae | *Nymphon brevirostre* |  |  |  | (3) third instar | flanking the midgut | LM, TEM, SEM | (Alexeeva et al., 2018) |
| Pantopoda | Nymphonoidea | Callipallenidae | *Propallene longiceps* |  |  |  | (3) third instar | flanking the midgut | LM | (Miyazaki and Makioka, 2012) |
| Xiphosura |  | Limulidae | *Limulus polyphemus* | Atlantic horseshoe crab |  |  | (3) segmented germ band | dorsal side of coelomic cavities | LM | (Kingsley, 1893) |
| Trombidiformes |  | Cheyletidae | *Cheyletus eruditus* |  |  |  | (3) segmented germ band | ventral mesoderm | LM | (Hafiz, 1935) |
| Trombidiformes |  | Tetranychidae | *Tetranychus urticae* | red spider mite, two-spotted spider mite | OB |  | (1) blastoderm | randomly distributed in yolk | LM, MM (*vasa in situ*) | (Dearden et al., 2003) |
| Sarcoptiformes |  | Acaridae | *Acarus siro* | flour mite |  |  | (3) segmented germ band | behind third larval leg | LM | (Hughes, 1950) |
| Opiliones | Eupnoi | Phalangiidae | *Opilio parietinus* |  |  |  | (2) early germ band | posterior germ band/blastopore | LM | (Holm, 1947) |
| Opiliones | Eupnoi | Phalangiidae | *Phalangium opilio* |  |  |  | (2) early germ band | posterior germ band/blastopore | LM, MM (*vasa in situ*) | (Faussek, 1888; Faussek, 1892; Gainett et al., 2022) |
| Scorpiones |  | Euscorpiidae | *Euscorpius carpathicus* |  |  |  | (2) early germ band | posterior germ disc/blastopore | LM | (Brauer, 1894) |
| Pseudoscorpiones |  | Chernetidae | *Pselaphochernes scorpioides* |  |  |  | (3) segmented germ band | mesoderm surrounding posterior midgut | LM | (Weygoldt, 1964) |
| Amblypygi |  | Phrynidae | *Heterophrynus pumilio* |  |  |  | (3) segmented germ band | ventral walls of mesodermal coelomic sacs | LM | (Gough, 1902) |
| Amblypygi |  | Phrynidae | *Phrynus marginemaculatus* | spotted tailless whip scorpion |  |  | (3) segmented germ band | ventral walls of mesodermal coelomic sacs | LM | (Weygoldt, 1975) |
| Uropygi |  | Thelyphonidae | *Thelyphonus caudatus* |  |  |  | (3) segmented germ band | ventral walls of mesodermal coelomic sacs | LM | (Schimkewitsch, 1903) |
| Araneae | Araneomorphae | Theridiidae | *Parasteatoda tepidariorum* | common house spider |  |  | (3) segmented germ band | ventral walls of mesodermal coelomic sacs | LM, MM (*vasa, piwi in situ*) | (Schwager et al., 2015) |
| Araneae | Araneomorphae | Agelenidae | *Agelena labyrinthica* |  |  |  | (3) segmented germ band | ventral walls of mesodermal coelomic sacs | LM | (Kautzsch, 1910) |
| **MYRIAPODA** | | | | | | | | | | |
| Scolopendromorpha |  | Scolopendridae | *Scolopendra cingulata* | Mediterranean banded centipede |  |  | (2) gastrulation | blastopore | LM | (Heymons, 1901) |
| Geophilomorpha |  | Linotaeniidae | *Strigamia maritima* |  |  |  | (2) blastoderm | blastopore | LM, MM (*vasa* and *nanos in situ*) | (Green and Akam, 2014) |
| Tetramerocerata |  | Pauropodidae | *Pauropus silvaticus* |  |  |  | (3) germ band | mesoderm | LM | (Tiegs, 1947) |
| Cephalostigmata |  | Scutigerellidae | *Hanseniella agilis* |  |  |  | (3) germ band | mesoderm | LM | (Tiegs, 1940) |
| Glomerida |  | Glomeridae | *Glomeris marginata* | pill millipede |  |  | (3) germ band | mesoderm in what will become the seventh and eighth leg segments as they form | LM | (Dohle, 1964) |
| **NON-HEXAPOD PANCRUSTACEA** | | | | | | | | | | |
| Podocopa |  | Cytherideidae | *Cyprideis torosa* |  |  |  | (2) gastrulation | from the mesoderm shortly after gastrulation | LM | (Weygoldt, 1960) |
| Calanoida |  | Calanidae | *Calanus finmarchicus* |  |  |  | (3) nauplius hatchling | from the mesoderm on either side of the intestine at posterior end of nauplius | LM | (Grobben, 1881) |
| Calanoida |  | Diaptomidae | *Eudiaptomus vulgaris* |  |  |  | (3) nauplius hatchling | from the mesoderm on either side of the intestine at posterior end of nauplius | LM | (Grobben, 1881) |
| Siphonostomatoida |  | Dichelesthiidae | unnamed species |  |  |  | (1) 32-cell stage | blastopore; last cell to bud out from the yolk and first cell to enter during gastrulation | LM | (McClendon, 1907) |
| Siphonostomatoida |  | Pandaridae | *Orthagoriscicola muricata* |  |  |  | (1) 32-cell stage | blastopore; last cell to bud out from the yolk and first cell to enter during gastrulation | LM | (McClendon, 1907) |
| Siphonostomatoida |  | Pandaridae | *Pandarus sinuatus* |  |  |  | (1) 32-cell stage | blastopore; last cell to bud out from the yolk and first cell to enter during gastrulation | LM | (McClendon, 1907) |
| Harpacticoida |  | Tisbidae | *Tisbe furcata* |  |  |  | (1) 64-cell stage | blastopore; last cell to bud out from the yolk and first cell to enter during gastrulation | LM | (Witschi, 1934) |
| Cyclopoida |  | Notodelphyidae | *Pachypygus gibber* |  |  |  | (2) gastrulation | blastopore, internalized soon after endoderm | LM | (Schimkewitsch, 1896) |
| Cyclopoida |  | Notodelphyidae | *Notopterophorus papilio* |  |  |  | (2) gastrulation | blastopore, internalized soon after endoderm | LM | (Schimkewitsch, 1896) |
| Leptostraca | Nebaliacea | Nebaliidae | *Nebalia bipes* |  |  |  | (3) late germ band | ventral walls of coelomic sacs | LM | (Manton, 1934) |
| Mysida |  | Mysidae | *Hemimysis lamornae* |  |  |  | (2) before gastrulation | center of germ disc/blastopore | LM | (Manton, 1928) |
| Mysida |  | Mysidae | *Neomysis integer* |  |  |  | (2) before gastrulation | center of germ disc/blastopore | LM | (Needham, 1937) |
| Mysida |  | Mysidae | *Mesopodopsis orientalis* |  |  |  | (2) before gastrulation | along median line of germ disc behind ectoteloblasts | LM | (Nair, 1939) |
| Tanaidacea |  | Leptocheliidae | *Heterotanais oerstedii* |  |  |  | (2) gastrulation | blastopore, internalized soon after yolk cells | LM | (Scholl, 1963) |
| Isopoda | Cymothoida | Bopyridae | *Bopyroides hippolytes* |  |  |  | (3) early germ band | lateral clusters, associated with liver rudiment | LM | (Strömberg, 1971) |
| Isopoda | Cymothoida | Anthuridae | *Cyathura polita* |  |  |  | (3) early germ band | lateral clusters, associated with liver rudiment | LM | (Strömberg, 1972) |
| Isopoda | Oniscidea | Ligiidae | *Ligia exotica* | sea roach |  |  | (3) just before hatching | last thoracic segment and first two pleonic segments | LM | (Terao and Cheng, 1926) |
| Isopoda | Oniscidea | Armadillidiidae | *Armadillidium nasatum* |  |  |  | (2) gastrulation | mesendodermal cluster at beginning of gastrulation | LM | (Goodrich, 1939) |
| Isopoda | Oniscidea | Armadillidiidae | *Armadillidium vulgare* | common pill-bug |  |  | (2) gastrulation | mesendodermal cluster at beginning of gastrulation | LM | (Lane, 1977) |
| Isopoda | Oniscidea | Porcellionidae | *Porcellio laevis* | swift woodlouse |  |  | (2) gastrulation | mesendodermal cluster at beginning of gastrulation | LM | (Goodrich 1939) |
| Isopoda | Oniscidea | Porcellionidae | *Porcellionides pruinosus* |  |  |  | (2) gastrulation | mesendodermal cluster at beginning of gastrulation | LM | (Lane, 1977) |
| Isopoda | Oniscidea | Porcellionidae | *Porcellio scaber* | common rough woodlouse |  |  | (2) gastrulation | mesendodermal cluster at beginning of gastrulation | LM | (Wolff, 2009) |
| Isopoda | Valvifera | Idoteidae | *Idotea granulosa* |  |  |  | (2) gastrulation | mesendodermal cluster at beginning of gastrulation | LM | (Strömberg, 1965) |
| Isopoda | Valvifera | Idoteidae | *Idotea neglecta* |  |  |  | (2) gastrulation | mesendodermal cluster at beginning of gastrulation | LM | (Strömberg, 1965) |
| Isopoda | Limnoriidea | Limnoriidae | *Limnoria lignorum* |  |  |  | (2) gastrulation | mesendodermal cluster at beginning of gastrulation | LM | (Strömberg, 1968) |
| Isopoda | Asellota | Asellidae | *Asellus aquaticus* | pond slater, water louse |  |  | (2) gastrulation | blastopore | LM | (Needham, 1942) |
| Amphipoda | Senticaudata | Gammaridae | *Gammarus pulex* |  |  |  | (2) gastrulation | mesendodermal cluster at beginning of gastrulation | LM | (Scholtz, 1990) |
| Amphipoda | Senticaudata | Talitridae | *Orchestia cavimana* |  |  |  | (1) 8-cell stage | *a* micromere | LM, CL | (Scholtz and Wolff, 2002; Wolff and Scholtz, 2002) |
| Amphipoda | Senticaudata | Hyalidae | *Parhyale hawaiensis* | sand flea | EX | EX | (1) 8-cell stage | *g* micromere (note: evidence that germ line can also regenerate) | LM, MM (*vasa*, *gcl*, *orb* *in situ*; Vasa antibody), EX (cell ablation) | (Extavour, 2005; Gupta and Extavour, 2013; Kaczmarczyk, 2014; Özhan-Kizil et al., 2009) |
| Anaspidacea |  | Anaspididae | *Anaspides tasmaniae* | mountain shrimp |  |  | (3) shortly before hatching | ventral walls of mesodermal coelomic sacs in T1/T2 | LM | (Hickman, 1936) |
| Euphausiacea |  | Euphausiidae | *Meganyctiphanes norvegica* | northern krill | OB |  | (1) 32-cell stage | X_d_ and X_v_ cells, sister to endoderm | LM, CL | (Alwes and Scholtz, 2004; Taube, 1909; Taube, 1915) |
| Decapoda | Dendrobranchiata | Sicyoniidae | *Sicyonia ingentis* | ridgeback prawn | OB |  | (2) gastrulation | MEvpp cell (medial ventral mesoderm) | LM, CL | (Hertzler, 2002; Pawlak et al., 2010) |
| Decapoda | Dendrobranchiata | Penaeidae | *Penaeus vannamei* | Pacific white shrimp, whiteleg shrimp | OB |  | (2) gastrulation | X_v_pp cell (ventral posterior mesendoderm) | LM, CL | (Hertzler, 2005; Pawlak et al., 2010) |
| Decapoda | Dendrobranchiata | Penaeidae | *Penaeus monodon* | black tiger shrimp | OB |  | (2) gastrulation | X_v_pp cell (ventral posterior mesendoderm) | LM, CL | (Biffis et al., 2009) |
| Decapoda | Dendrobranchiata | Penaeidae | *Penaeus japonicus* | Kuruma shrimp; Japanese tiger shrimp | OB |  | (2) gastrulation | X_v_pp cell (ventral posterior mesendoderm) | LM, CL, TEM | (Grattan et al., 2013; Pawlak et al., 2010; Vincent and Hertzler, 2018) |
| Decapoda | Pleocyemata (Caridea) | Atyidae | *Caridina laevis* |  |  |  | (3) segmented embryo | ventral mesoderm on either side of the gut | LM | (Nair, 1949) |
| Decapoda | Pleocyemata (Caridea) | Palaemonidae | *Macrobrachium nipponense* | oriental river prawn | OB |  | (1) 16-cell stage | unnamed blastomere | LM, MM (*vasa in situ*, Vasa antibody), TEM | (Chen et al., 2021; Ma et al., 2019; Qiu et al., 2013) |
| Decapoda | Pleocyemata (Reptantia, Achelata) | Palinuridae | *Panulirus japonicus* | Japanese spiny lobster |  |  | (3) segmented embryo | ventral walls of mesodermal coelomic sacs in T1 | LM | (Terao, 1929) |
| Anostraca | Artemiina | Artemiidae | *Artemia salina* |  |  |  | (3) nauplius hatchling | mesoderm of first trunk segment | LM | (Anderson, 1967) |
| Notostraca |  | Triopsidae | *Triops longicaudatus* |  |  |  | (3) fourth instar larva | on either side of the gut | LM | (Mitsumoto and Makioka, 2002) |
| Notostraca |  | Triopsidae | *Triops granarius* |  |  |  | (3) fourth instar larva | on either side of the gut | LM | (Mitsumoto and Makioka, 2003) |
| Spinicaudata |  | Limnadiidae | *Paralimnadia stanleyana* |  |  |  | (3) nauplius hatchling | mesoderm of first trunk segment | LM | (Anderson, 1967) |
| Ctenopoda |  | Holopediidae | *Holopedium gibberum* |  |  |  | (1) 16-cell stage | D^IV 1^ blastomere at the vegetal pole, which becomes the site of gastrulation | LM | (Baldass, 1937) |
| Ctenopoda |  | Sididae | *Penilia avirostris* |  |  |  | (3) late embryo | mesoderm on either side of intestine | LM | (Sudler, 1899) |
| Anomopoda |  | Macrothiricidae | *Lathonura rectirostris* |  |  |  | (2) gastrulation | four cells at blastopore lip | LM | (Grobben, 1879) |
| Anomopoda |  | Daphniidae | *Simocephalus vetulus* |  |  |  | (2) late blastoderm | posterior “ventral mass” on inside of blastoderm, later the site of gastrulation | LM | (Cannon, 1921) |
| Anomopoda |  | Daphniidae | *Daphnia magna* |  | OB |  | (1) 16-cell stage | unnamed blastomere | LM, MM (Vasa antibody) | (Sagawa et al., 2005) |
| Onychopoda |  | Cercopagididae | *Bythotrephes longimanus* | spiny water flea | OB |  | (1) 16-cell stage | one daughter cell of the dI micromere | LM | (Alwes and Scholtz, 2014) |
| Onychopoda |  | Polyphemidae | *Polyphemus pediculus* |  | OB |  | (1) 16-cell stage | one daughter cell of the dI micromere | LM, CL | (Kühn, 1913) |
| **HEXAPODA** | | | | | | | | | | |
| ***Ametabola*** | | | | | | | | | | |
| Poduromorpha |  | Neanuridae | *Anurida maritima* | seashore springtail |  |  | (3) segmented germ band | abdominal mesoderm | LM | (Claypole, 1898) |
| Poduromorpha |  | Onychiuridae | *Tetrodontophora bielanensis* | giant springtail | OB |  | (1) cleavage stage embryo | region of cortical cytoplasm in early-stage embryo | TEM | (Klag, 1982) |
| Symphypleona |  | Sminthuridae | *Allacma fusca* |  |  |  | (1) blastoderm-stage embryo | scattered in yolk | TEM | (Klag and Światek, 1999) |
| Entomobryomorpha |  | Tomoceridae | *Tomocerus cuspidatus* |  |  |  | (1) cleavage-stage embryo | scattered in the yolk, inner mass | LM, TEM | (Tomizuka and Machida, 2015) |
| Entomobryomorpha |  | Tomoceridae | *Tomocerus ishibashii* |  |  |  | (1) blastoderm-stage embryo | center of yolk after blastoderm formation | LM | (Uemiya and Ando, 1991) |
| Archaeognatha |  | Machilidae | *Petrobius brevistylis* |  |  |  | (3) segmented germ band | dorsal coelomic sacs of T2–A1 mesoderm | LM | (Larink, 1969) |
| Zygentoma |  | Lepismatidae | *Lepisma saccharina* | silverfish |  |  | (2) early germ band | posterior end of germ band; author notes difficulty in following migration of putative germ cells | LM | (Heymons, 1897) |
| Zygentoma |  | Lepismatidae | *Thermobia domestica* | firebrat |  |  | (3) segmented germ band | dorsal coelomic sacs of abdominal mesoderm | LM | (Woodland, 1957) |
| ***Palaeoptera*** | | | | | | | | | | |
| Odonata | Epiprocta | Epiophlebiidae | *Epiophlebia superstes* |  |  |  | (2) cellular blastoderm | posterior blastoderm; unclear whether these cells are the same cells that appear in the gonads later in embryogenesis | LM | (Ando, 1962) |
| ***Polyneoptera*** | | | | | | | | | | |
| Dermaptera |  | Forficulidae | *Forficula auricularia* | common earwig |  |  | (2) cellular blastoderm | posterior blastoderm | LM | (Heymons, 1895) |
| Dermaptera |  | Labiduridae | *Labidura riparia* | shore earwig | OB |  | (1) syncytial blastoderm | posterior blastoderm | LM | (Singh, 1967) |
| Orthoptera | Ensifera | Gryllidae | *Oecanthus niveus* | narrow-winged tree cricket |  |  | (3) late germ band | on either side of the dorsal vessel | LM | (Ayers, 1884) |
| Orthoptera | Ensifera | Gryllidae | *Acheta domesticus* | house cricket |  |  | (2) early germ band | posterior end of germ band | LM | (Heymons, 1895) |
| Orthoptera | Ensifera | Gryllidae | *Gryllus campestris* | European field cricket |  |  | (2) early germ band | posterior end of germ band | LM | (Heymons, 1895) |
| Orthoptera | Ensifera | Gryllidae | *Gryllus bimaculatus* | two-spotted field cricket |  | EX | (3) segmented germ band | abdominal mesoderm, before formation of coelomic sacs | LM, MM (Vasa and Piwi antibodies, many *in situ*), EX (genetic manipulation) | (Donoughe et al., 2014; Ewen-Campen et al., 2013a; Nakamura and Extavour, 2016) |
| Orthoptera | Ensifera | Tettigoniidae | *Conocephalus brevipennis* | short-winged meadow katydid |  |  | (3) late germ band | inner walls of coelomic sacs in A1–A6 mesoderm | LM | (Wheeler, 1893) |
| Orthoptera | Caelifera | Acrididae | *Melanoplus differentialis* | differential grasshopper |  |  | (2) early germ band | lateral margins of germ band, A1–A9 | LM | (Nelsen, 1934) |
| Orthoptera | Caelifera | Acrididae | *Locusta migratoria* | migratory locust |  |  | (3) late germ band | dorsal coelomic sacs of A2–A5 mesoderm | LM | (Roonwal, 1937) |
| Orthoptera | Caelifera | Acrididae | *Schistocerca gregaria* | desert locust |  |  | (2) early germ band | lateral margins of abdominal germ band | LM, MM (Vasa antibody) | (Chang et al., 2002) |
| Blattodea | Solumblattodea | Blattidae | *Periplaneta orientalis* | oriental cockroach |  |  | (2) early germ band | posterior end of germ band | LM | (Heymons, 1895) |
| Blattodea | Blaberoidea | Ectobiidae | *Blattella germanica* | German cockroach |  |  | (2) early germ band | posterior end of germ band; author notes that putative germ cells are indistinguishable from mesodermal cells but identifies them as germ cells based on comparison with *Periplaneta orientalis* | LM | (Heymons, 1895) |
| Phasmatodea | Oriophasmata | Lonchodidae | *Carausius morosus* | common stick insect |  |  | (2) early germ band or (3) late germ band | (Hammerschmidt, 1910; Wiesmann, 1926) say posterior end of germ band; (Cavallin, 1970) says abdominal mesoderm | LM | (Cavallin, 1970; Hammerschmidt, 1910; Wiesmann, 1926) |
| Mantodea |  | Mantidae | *Rhombodera crassa* |  |  |  | (3) late germ band | coelomic sacs in A3–A5 mesoderm | LM | (Görg, 1959) |
| ***Condylognatha + Psocodea*** | | | | | | | | | | |
| Thysanoptera | Tubulifera | Phlaeothripidae | *Haplothrips verbasci* |  | OB |  | (1) syncytial blastoderm | posterior blastoderm | LM | (Heming, 1979) |
| Thysanoptera | Tubulifera | Phlaeothripidae | *Bactrothrips brevitubus* |  | OB |  | (1) syncytial blastoderm | posterior blastoderm | LM | (Haga, 1985) |
| Hemiptera | Heteroptera (Pentatomomorpha) | Lygaeidae | *Oncopeltus fasciatus* | large milkweed bug |  |  | (2) cellular blastoderm | posterior blastoderm | LM, MM (Vasa antibody, many *in situ* hybridization) | (Butt, 1949; Ewen-Campen et al., 2013b; Kao et al., 2025) |
| Hemiptera | Heteroptera (Pentatomomorpha) | Blissidae | *Blissus leucopterus hirtus* | hairy chinch bug |  |  | (2) cellular blastoderm | posterior blastoderm | LM | (Choban and Gupta, 1972) |
| Hemiptera | Heteroptera (Pentatomomorpha) | Pyrrhocoridae | *Pyrrhochoris apterus* | European firebug |  |  | (3) segmented germ band | from the mesoderm in segments A1–A8 | LM | (Seidel, 1924) |
| Hemiptera | Heteroptera (Cimicomorpha) | Reduviidae | *Rhodnius prolixus* | kissing bug |  |  | (2) cellular blastoderm | posterior blastoderm | LM, SEM | (Heming and Huebner, 1994; Kelly and Huebner, 1989; Mellanby, 1935) |
| Hemiptera | Sternorrhynca | Monophlebidae | *Icerya purchasi* | cottony cushion scale |  |  | (2) cellular blastoderm | posterior blastoderm | LM | (Shinji, 1919) |
| Hemiptera | Sternorrhynca | Pseudococcidae | *Pseudococcus citri* | citrus mealybug |  |  | (2) cellular blastoderm | posterior blastoderm | LM | (Schrader, 1922) |
| Hemiptera | Sternorrhynca | Aphididae | *Acyrthosiphon pisum* | pea aphid | OB |  | (2) cellular blastoderm | posterior blastoderm | LM, MM (Vasa, Nanos antibody, *vasa in situ*) | (Chang et al., 2006; Chang et al., 2007; Lin et al., 2014; Miura et al., 2003) |
| Psocodea | Psocomorpha | Archipsocidae | *Archipsocus fernandi* |  |  |  | (2) cellular blastoderm | posterior blastoderm | LM | (Fernando, 1934) |
| Psocodea | Troctomorpha | Liposcelididae | *Liposcelis divergens* | book louse |  |  | (2) cellular blastoderm | posterior blastoderm | LM | (Goss, 1952; Goss, 1953) |
| ***Holometabola*** | | | | | | | | | | |
| Hymenoptera | Apocrita (Aculeata) | Formicidae | *Lasius niger* | black garden ant | OB |  | (1) cellular blastoderm | posterior pole of blastoderm | LM, MM (Vasa antibody, *nanos, oskar in situ*) | (Rafiqi et al., 2020) |
| Hymenoptera | Apocrita (Aculeata) | Formicidae | *Camponotus floridanus* | Florida carpenter ant | OB |  | (1) cellular blastoderm | posterior pole of blastoderm | LM, MM (Vasa antibody, *nanos, oskar in situ*) | (Rafiqi et al., 2020) |
| Hymenoptera | Apocrita (Aculeata) | Formicidae | *Messor pergandei* | black harvester ant | OB |  | (1) cellular blastoderm | posterior pole of blastoderm | LM, MM (*oskar, vasa, nanos in situ*) | (Khila and Abouheif, 2010; Lynch et al., 2011) |
| Hymenoptera | Apocrita (Aculeata) | Apidae | *Apis mellifera* | western honeybee |  |  | (3) segmented germ band | genital ridges in A3–A6 mesoderm | LM, SEM, MM (*vasa, nanos in situ*) | (Dearden, 2006; Fleig and Sander, 1986; Nelson, 1915) |
| Hymenoptera | Apocrita (Parasitoida) | Pteromalidae | *Nasonia vitripennis* | jewel wasp | EX |  | (1) syncytial blastoderm (nuclear cycle 8) | pole cells at posterior pole of blastoderm | LM, MM (*nanos, oskar in situ*), EX (genetic manipulation), TS | (Bull, 1982; Lynch and Desplan, 2010; Lynch et al., 2011; Quan et al., 2019) |
| Hymenoptera | Apocrita (Parasitoida) | Trichogrammatidae | *Trichogramma evanescens* |  | OB |  | (1) syncytial blastoderm | pole cells at posterior pole of blastoderm | LM | (Gatenby, 1917; Gatenby, 1918) |
| Hymenoptera | Apocrita (Parasitoida) | Encyrtidae | *Copidosoma floridanum* |  | EX |  | (1) 4-cell stage | B4 blastomere | LM, MM (Vasa *in situ* and antibody), EX (cell ablation) | (Donnell et al., 2004; Grbic’, 2003; Zhurov et al., 2004) |
| Hymenoptera | Apocrita (Parasitoida) | Braconidae | *Macrocentrus cingulum* |  | OB |  | (2) cellular blastoderm | inside the posterior blastocoel | LM, MM (Vasa antibody) | (Grbic’, 2003; Sucena et al., 2014) |
| Hymenoptera | Apocrita (Parasitoida) | Braconidae | *Coeloides vancouverensis* |  | OB |  | (1) syncytial blastoderm | pole cells at posterior pole of blastoderm | LM | (Ryan, 1963) |
| Hymenoptera | Apocrita (Parasitoida) | Braconidae | *Habrobracon juglandis* |  | OB |  | (1) syncytial blastoderm | pole cells at posterior pole of blastoderm | LM | (Amy, 1961) |
| Hymenoptera | Apocrita (Parasitoida) | Ichneumonidae | *Cosmoconus meridionator* |  | OB |  | (1) syncytial blastoderm | pole cells at posterior pole of blastoderm | TEM | (Klag and Bilinski, 1993) |
| Hymenoptera | Apocrita (Parasitoida) | Ichneumonidae | *Lissonota catenator* |  | OB |  | (1) syncytial blastoderm | pole cells at posterior pole of blastoderm | TEM | (Klag and Bilinski, 1993) |
| Hymenoptera | Apocrita (Parasitoida) | Ichneumonidae | *Pimpla turionellae* |  | EX | EX | (1) syncytial blastoderm | pole cells at posterior pole of blastoderm (note: evidence that germ line can also regenerate) | LM, EX (damage cytoplasm) | (Achtelig and Krause, 1971; Bronskill, 1959; Meng, 1968) |
| Hymenoptera | Eusymphyta | Tenthredinidae | *Nematus ribesii* | gooseberry sawfly |  |  | (3) segmented germ band | posterior midgut rudiment | LM | (Shafiq, 1954) |
| Hymenoptera | Eusymphyta | Tenthredinidae | *Athalia rosae* | turnip sawfly |  |  | (1) syncytial blastoderm | pole cells at posterior pole of blastoderm | LM, MM (Vasa antibody) | (Nakao et al., 2006) |
| Raphidioptera |  | Inocelliidae | *Inocellia japonica* |  |  |  | (1) syncytial blastoderm | pole cells at posterior pole of blastoderm | LM, SEM | (Tsutsumi and Machida, 2006) |
| Megaloptera |  | Sialidae | *Sialis mitsuhashii okamoto* | alderfly |  |  | (2) late blastoderm | posterior end of ventral plate | TEM | (Suzuki et al., 1981) |
| Strepsiptera |  | Stylopidae | *Stylops spp.* |  |  |  | (3) extended germ band | posterior end of the germ band | LM | (Noskiewicz and Poluszyński, 1927) |
| Coleoptera | Adephaga | Carabidae | *Carabus insulicola* |  |  |  | ?? | no pole cells observed | LM, SEM | (Kobayashi et al., 2013) |
| Coleoptera | Adephaga | Gyrinidae | *Dineutus mellyi* | whirligig beetle |  |  | ?? | no pole cells observed | LM, SEM | (Komatsu and Kobayashi, 2012) |
| Coleoptera | Adephaga | Dytiscidae | *Hydaticus pacificus* | diving beetle |  |  | ?? | no pole cells observed | LM, SEM | (Niikura et al., 2017) |
| Coleoptera | Polyphaga (Elateroidea) | Lampyridae | *Nipponoluciola cruciata* |  |  |  | ?? | no pole cells observed | LM | (Kobayashi and Ando, 1985) |
| Coleoptera | Polyphaga (Elateroidea) | Lampyridae | *Pyrocoelia rufa* |  |  |  | ?? | no pole cells observed | LM | (Kobayashi et al., 2006) |
| Coleoptera | Polyphaga (Hydrophiloidea) | Hydrophilidae | *Hydrophilus piceus* | great silver water beetle |  |  | (3) segmented germ band | from the walls of the coelomic sacs of abdominal mesoderm | LM | (Heider, 1889) |
| Coleoptera | Polyphaga (Scarabaeoidea) | Passalidae | *Odontotaenius disjunctus* | patent-leather beetle, horned passalus |  |  | (3) late germ band | dorsal side of abdominal germ band, near A6–A7 | LM | (Krause, 1947) |
| Coleoptera | Polyphaga (Scarabaeoidea) | Scarabaeidae | *Phyllophaga fervida* |  |  |  | (2) cellular blastoderm | posterior blastoderm | LM | (Luginbill, 1953) |
| Coleoptera | Polyphaga (Bostrichoidea) | Dermestidae | *Dermestes frischii* |  | OB |  | (2) cellular blastoderm | posterior blastoderm | LM | (Küthe, 1966) |
| Coleoptera | Polyphaga (Tenebrionoidea) | Meloidae | *Lytta viridana* | blister beetle |  |  | (2) cellular blastoderm | posterior blastoderm | LM | (Church and Rempel, 1971; Rempel and Church, 1969) |
| Coleoptera | Polyphaga (Tenebrionoidea) | Tenebrionidae | *Tenebrio molitor* | yellow mealworm beetle |  |  | (2) cellular blastoderm | posterior blastoderm | LM | (Ullmann, 1964) |
| Coleoptera | Polyphaga (Tenebrionoidea) | Tenebrionidae | *Tribolium confusum* | confused flour beetle |  |  | (2) cellular blastoderm | posterior blastoderm | LM | (Hodson, 1934; Stanley and Grundmann, 1970) |
| Coleoptera | Polyphaga (Tenebrionoidea) | Tenebrionidae | *Tribolium castaneum* | red flour beetle |  | EX | (2) cellular blastoderm | posterior blastoderm | LM, MM (*vasa in situ*), EX (genetic manipulation) | (Ansari et al., 2018; Schröder, 2006) |
| Coleoptera | Polyphaga (Curculionidea) | Curculionidae | *Sphenophorus callosus* | southern corn billbug | EX |  | (1) syncytial blastoderm | pole cells at posterior pole of blastoderm | LM, EX (cytoplasmic cauterization) | (Hegner, 1911; Hegner, 1914; Wray, 1937) |
| Coleoptera | Polyphaga (Curculionidea) | Curculionidae | *Otiorhynchus ligustici* | alfalfa snout beetle | OB |  | (1) syncytial blastoderm | pole cells at posterior pole of blastoderm | LM | (Butt, 1936) |
| Coleoptera | Polyphaga (Curculionidea) | Curculionidae | *Sitophilus oryzae* | rice weevil |  |  | (1) syncytial blastoderm | pole cells at posterior pole of blastoderm | LM | (Tiegs and Murray, 1938) |
| Coleoptera | Polyphaga (Chrysomeloidea) | Chrysomelidae | *Calligrapha multipunctata* | common willow calligrapha | EX |  | (1) syncytial blastoderm | pole cells at posterior pole of blastoderm | LM, EX (cytoplasmic removal) | (Hegner, 1908; Hegner, 1909a; Hegner, 1909b) |
| Coleoptera | Polyphaga (Chrysomeloidea) | Chrysomelidae | *Leptinotarsa decemlineata* | Colorado potato beetle | EX |  | (1) syncytial blastoderm | pole cells at posterior pole of blastoderm | LM, EX (cytoplasmic cauterization/removal) | (Hegner, 1908; Hegner, 1911; Hegner, 1914) |
| Coleoptera | Polyphaga (Chrysomeloidea) | Chrysomelidae | *Callosobruchus maculatus* | cowpea weevil | EX |  | (1) syncytial blastoderm | pole cells at posterior pole of blastoderm | LM, MM (*oskar, vasa, tudor in situ*), EX (UV irradiation) | (Brauer, 1925; Brauer, 1949; Quan, 2018) |
| Coleoptera | Polyphaga (Chrysomeloidea) | Chrysomelidae | *Acanthoscelides obtectus* | bean weevil | OB |  | (1) syncytial blastoderm (nuclear cycle 8) | pole cells at posterior pole of blastoderm | LM, MM (*vasa in situ*), TEM | (Jung, 1966; Lynch et al., 2011) |
| Coleoptera | Polyphaga (Chrysomeloidea) | Chrysomelidae | *Euryope terminalis* | milkweed leaf beetle | OB |  | (1) syncytial blastoderm | pole cells at posterior pole of blastoderm | LM | (Paterson, 1931) |
| Coleoptera | Polyphaga (Chrysomeloidea) | Chrysomelidae | *Platycorynus compressicornis* |  | OB |  | (1) syncytial blastoderm | pole cells at posterior pole of blastoderm | LM | (Paterson, 1935) |
| Trichoptera | Annulipalpia | Stenopsychidae | *Stenopsyche griseipennis* |  |  |  | (3) segmented germ band | genital ridges in A2–A7 mesoderm | LM | (Miyakawa, 1974) |
| Lepidoptera | Micropterigoidea | Micropterigidae | *Neomicropteryx nipponensis* |  |  |  | (3) segmented germ band | mediodorsal coelomic sacs in A5 mesoderm | LM | (Kobayashi and Ando, 1984) |
| Lepidoptera | Glossata (Hepialoidea) | Hepialidae | *Endoclita signifer* |  |  |  | (2) early germ band | posterior germ band | LM | (Ando and Tanaka, 1980) |
| Lepidoptera | Apoditrysia (Tortrichoidea) | Tortricidae | *Epiphyas postvittana* | light brown apple moth |  |  | (2) early germ band | between germ band and yolk | LM | (Anderson and Wood, 1968) |
| Lepidoptera | Obtectomera (Papilionoidea) | Nymphalidae | *Euvanessa (Nymphalis) antiopa* | mourning cloak |  |  | (2) early germ band | midline of ventral plate, about one-third length from the posterior | LM | (Woodworth, 1889) |
| Lepidoptera | Obtectomera (Papilionoidea) | Nymphalidae | *Pararge aegeria* | speckled wood butterfly | OB |  | (2) cellular blastoderm | ventral blastoderm | LM, MM (*nanos in situ*), TS | (Carter et al., 2013; Carter et al., 2015) |
| Lepidoptera | Obtectomera (Papilionoidea) | Pieridae | *Pieris rapae* | cabbage white, cabbage butterfly |  |  | (3) segmented germ band | coelomic sacs of A4–A5 mesoderm | LM | (Eastham, 1931) |
| Lepidoptera | Obtectomera (Papilionoidea) | Papilionidae | *Parnassius glacialis* | glacial Apollo, Japanese clouded Apollo |  |  | (2) late cellular blastoderm/early germ band | midline of ventral plate, about one-third length from the posterior | LM | (Tanaka, 1987) |
| Lepidoptera | Obtectomera (Papilionoidea) | Papilionidae | *Luehdorfia japonica* | Gifu butterfly |  |  | (2) early germ band | midline of germ band, about one-third length from the posterior | LM | (Tanaka, 1987) |
| Lepidoptera | Obtectomera (Papilionoidea) | Papilionidae | *Byasa alcinous alcinous* | Chinese windmill |  |  | (2) early germ band | midline of germ band, about one-third length from the posterior | LM | (Tanaka, 1987) |
| Lepidoptera | Obtectomera (Gelechioidea) | Gelechiidae | *Pectinophora gossypiella* | pink bollworm | OB |  | (2) cellular blastoderm | posterior, ventral blastoderm | TEM | (Berg and Gassner, 1978) |
| Lepidoptera | Macroheterocera (Noctuoidea) | Amatidae | *Amata fortunei* | white-spotted moth |  |  | (2) early germ band | midline of germ band, near the center | LM | (Tanaka, 1985) |
| Lepidoptera | Macroheterocera (Noctuoidea) | Noctuidae | *Helicoverpa zea* | corn earworm |  |  | (2) early germ band | midline of germ band, near the center | LM | (Presser and Rutschky, 1957) |
| Lepidoptera | Macroheterocera (Noctuoidea) | Arctiinae | *Spilosoma virginica* | Virginian tiger moth |  |  | (3) late germ band | coelomic sacs in abdominal mesoderm | LM | (Johannsen, 1929) |
| Lepidoptera | Macroheterocera (Bombycoidea) | Bombycidae | *Bombyx mori* | domestic silk moth | EX | EX | (2) cellular blastoderm | midline of ventral plate, about one-third length from the posterior |  | (Miya, 1958; Nakao, 1999; Nakao and Takasu, 2019) |
| Lepidoptera | Macroheterocera (Bombycoidea) | Saturniidae | *Antheraea pernyi* | Chinese (oak) tussar moth |  |  | (3) segmented germ band | mesodermal coelomic sacs of all abdominal segments | LM | (Saito, 1937) |
| Mecoptera |  | Panorpidae | *Panorpa pryeri* |  | OB |  | (2) early germ band | posterior end of germ band | LM | (Suzuki, 1990) |
| Mecoptera |  | Panorpodidae | *Panorpodes paradoxa* |  |  |  | (2) early germ band | posterior end of germ band | LM | (Suzuki, 1990) |
| Mecoptera |  | Bittacidae | *Bittacus laevipes* |  |  |  | (2) early germ band | posterior end of germ band | LM | (Suzuki, 1990) |
| Mecoptera |  | Boreidae | *Boreus westwoodi* |  |  |  | (2) early germ band | posterior end of germ band | LM | (Suzuki, 1990) |
| Siphonaptera |  | Pulicidae | *Ctenocephalides felis* | cat flea |  |  | (1) syncytial blastoderm (nuclear cycle 6) | pole cells at posterior pole of blastoderm | LM | (Kessel, 1939) |
| Diptera | Psychodomorpha | Psychodidae | *Phlebotomus papatasi* | sand fly | OB |  | (1) syncytial blastoderm | pole cells at posterior pole of blastoderm | LM | (Abbassy et al., 1995a; Abbassy et al., 1995b; Abbassy et al., 1995c) |
| Diptera | Psychodomorpha | Psychodidae | *Clogmia albipunctata* | moth fly |  | EX | (2) cellular blastoderm | posterior blastoderm; no morphologically distinct pole cells | SEM, LM, MM (Vasa antibody, *nanos in situ*), EX (genetic manipulation) | (Jiménez-Guri et al., 2014; Yoon et al., 2019) |
| Diptera | Culicomorpha (Chironomoidea) | Simuliidae | *Simulium pictipes* | black fly | OB |  | (1) syncytial blastoderm | pole cells at posterior pole of blastoderm | LM | (Gambrell, 1933) |
| Diptera | Culicomorpha (Chironomoidea) | Chironomidae | *Chironomus sp.* |  |  |  | (1) syncytial blastoderm | pole cells at posterior pole of blastoderm | LM | (Klomp et al., 2015; Weissmann, 1863) |
| Diptera | Culicomorpha (Chironomoidea) | Chironomidae | *Chironomus confinis* | nonbiting midge | OB |  | (1) syncytial blastoderm | pole cells at posterior pole of blastoderm | LM | (Hegner, 1914) |
| Diptera | Culicomorpha (Chironomoidea) | Chironomidae | *Smittia sp.* |  | OB |  | (1) syncytial blastoderm | pole cells at posterior pole of blastoderm | LM, TEM | (Zissler and Sander, 1973) |
| Diptera | Culicomorpha (Culicoidea) | Culicidae | *Aedes aegypti* | yellow fever mosquito | OB |  | (1) syncytial blastoderm | pole cells at posterior pole of blastoderm | LM, MM (*oskar in situ*) | (Juhn and James, 2006; Raminani and Cupp, 1975) |
| Diptera | Culicomorpha (Culicoidea) | Culicidae | *Culex quinquefasciatus* | southern house mosquito | OB |  | (1) syncytial blastoderm | pole cells at posterior pole of blastoderm | LM, MM (*nanos*, *oskar in situ*) | (Davis, 1967; Juhn et al., 2008) |
| Diptera | Culicomorpha (Culicoidea) | Culicidae | *Culiseta inornata* | winter marsh mosquito | OB |  | (1) syncytial blastoderm | pole cells at posterior pole of blastoderm | LM | (Harber and Mutchmor, 1970) |
| Diptera | Culicomorpha (Culicoidea) | Culicidae | *Anopheles gambiae* | African malaria mosquito | OB |  | (1) syncytial blastoderm (nuclear cycle 9) | pole cells at posterior pole of blastoderm | LM, MM (*oskar in situ*) | (Juhn and James, 2006) |
| Diptera | Bibionomorpha | Cecidomyiidae | *Miastor metraloas* | gall midge | OB |  | (1) syncytial blastoderm (nuclear cycle 3) | pole cells at posterior pole of blastoderm | LM, TEM | (Hegner, 1914; Mahowald, 1975) |
| Diptera | Bibionomorpha | Cecidomyiidae | *Mayetiola destructor* | Hessian fly |  |  | (1) syncytial blastoderm (nuclear cycle 5) | pole cells at posterior pole of blastoderm | LM | (Metcalfe, 1935) |
| Diptera | Bibionomorpha | Sciaridae | *Bradysia coprophila* | black fungus gnat | OB |  | (1) syncytial blastoderm (nuclear cycle 5) | pole cells at posterior pole of blastoderm | LM | (DuBois, 1932; Phalle and Sullivan, 1996) |
| Diptera | Bibionomorpha | Sciaridae | *Sciara sp.* |  | OB |  | (1) syncytial blastoderm (nuclear cycle 5) | pole cells at posterior pole of blastoderm | LM | (Butt, 1934) |
| Diptera | Cyclorrapha (Platypezoidea) | Phoridae | *Megaselia abdita* | scuttle fly | OB |  | (1) syncytial blastoderm (nuclear cycle 9) | pole cells at posterior pole of blastoderm | SEM, LM, MM (Vasa antibody) | (Wotton et al., 2014) |
| Diptera | Schizophora (Tephritoidea) | Tephritidae | *Bactrocera tryoni* | Queensland fruit fly | OB |  | (1) syncytial blastoderm (nuclear cycle 10) | pole cells at posterior pole of blastoderm | LM | (Anderson, 1962) |
| Diptera | Schizophora (Ephydroidea) | Drosophilidae | *Drosophila melanogaster* | common fruit fly, vinegar fly | EX |  | (1) syncytial blastoderm (nuclear cycle 9) | pole cells at posterior pole of blastoderm | LM, EM, MM (many), EX (genetic manipulation, cytoplasmic transplant) | (Counce, 1963; Ephrussi and Lehmann, 1992; Huettner, 1923; Illmensee and Mahowald, 1974; Mahowald, 1968) |
| Diptera | Schizophora (Ephydroidea) | Drosophilidae | *Drosophila hydei* |  | OB |  | (1) syncytial blastoderm | pole cells at posterior pole of blastoderm | LM, TEM | (Counce, 1963; Mahowald, 1968) |
| Diptera | Schizophora (Ephydroidea) | Drosophilidae | *Drosophila willistoni* |  | OB |  | (1) syncytial blastoderm | pole cells at posterior pole of blastoderm | LM, TEM | (Counce, 1963; Mahowald, 1968) |
| Diptera | Schizophora (Ephydroidea) | Drosophilidae | *Drosophila immigrans* |  | EX |  | (1) syncytial blastoderm | pole cells at posterior pole of blastoderm | TEM, EX (cytoplasmic transplant) | (Mahowald, 1968; Mahowald et al., 1976) |
| Diptera | Schizophora (Ephydroidea) | Drosophilidae | *Drosophila busckii* |  | OB |  | (1) syncytial blastoderm | pole cells at posterior pole of blastoderm | LM | (Counce, 1963) |
| Diptera | Schizophora (Ephydroidea) | Drosophilidae | *Drosophila virilis* |  | OB |  | (1) syncytial blastoderm | pole cells at posterior pole of blastoderm | LM, MM (Vasa, Oskar antibody, many *in situ*) | (Counce, 1963; Rivard et al., 2025) |
| Diptera | Schizophora (Ephydroidea) | Drosophilidae | *Drosophila americana* |  | OB |  | (1) syncytial blastoderm | pole cells at posterior pole of blastoderm | LM | (Counce, 1963) |
| Diptera | Schizophora (Ephydroidea) | Drosophilidae | *Drosophila repleta* |  | OB |  | (1) syncytial blastoderm | pole cells at posterior pole of blastoderm | LM | (Counce, 1963) |
| Diptera | Schizophora (Ephydroidea) | Drosophilidae | *Drosophila gibberosa* |  | OB |  | (1) syncytial blastoderm | pole cells at posterior pole of blastoderm | LM | (Counce, 1963) |
| Diptera | Schizophora (Ephydroidea) | Drosophilidae | *Drosophila funebris* |  | OB |  | (1) syncytial blastoderm | pole cells at posterior pole of blastoderm | LM | (Counce, 1963) |
| Diptera | Schizophora | Hippoboscidae | *Melophagus ovinus* | sheep ked, sheep tick | OB |  | (1) syncytial blastoderm | pole cells at posterior pole of blastoderm | LM | (Lassmann, 1936) |
| Diptera | Schizophora | Muscidae | *Stomoxys calcitrans* | stable fly |  |  | (1) syncytial blastoderm | pole cells at posterior pole of blastoderm | LM, TEM, SEM | (Ajidagba et al., 1983) |
| Diptera | Schizophora | Muscidae | *Musca domestica* | house fly | OB |  | (1) syncytial blastoderm (nuclear cycle 6) | pole cells at posterior pole of blastoderm | LM | (West et al., 1968) |
| Diptera | Schizophora (Oestroidea) | Calliphoridae | *Calliphora vicina* | blowfly | OB |  | (1) syncytial blastoderm | pole cells at posterior pole of blastoderm | LM | (Hegner, 1914) |
| Diptera | Schizophora (Oestroidea) | Calliphoridae | *Phormia regina* | black blowfly | OB |  | (1) syncytial blastoderm | pole cells at posterior pole of blastoderm | LM | (Auten, 1934) |
| Diptera | Schizophora (Oestroidea) | Calliphoridae | *Lucilia sericata* | sheep blowfly | OB |  | (1) syncytial blastoderm (nuclear cycle 10) | pole cells at posterior pole of blastoderm | LM | (Davis, 1967; Fish, 1947a; Fish, 1947b; Fish, 1949; Fish, 1952) |

#### **Supplementary Table S2. Studies used for phylogram and constraint tree construction.** Clade: clade extracted from the cited study; Resolved rank: lowest taxonomic rank resolved in our phylogram from the study.

| **Reference** | **Clade** | **Resolved rank** | **Note** |
| --- | --- | --- | --- |
| (Howard et al., 2022) | Panarthropoda | Phylum |  |
| (Baker et al., 2021) | Onychophora | Family |  |
| (Giribet, 2018) | Chelicerata | Order | Euchelicerate relationships outside of Arachnopulmonata were left as a polytomy to reflect the continued uncertainty in chelicerate phylogeny (Giribet, 2018; Sharma and Gavish-Regev, 2025). |
| (Sabroux et al., 2023) | Pantopoda | Family |  |
| (Fernández et al., 2018) | Myriapoda | Class |  |
| (Bernot et al., 2023) | Pancrustacea | Family, Order, Subphylum | Lower-rank constraints for Hexapoda, Decapoda, Isopoda, Amphipoda, Diplostraca, and Copepoda were substituted from other studies. |
| (Wolfe et al., 2019) | Decapoda | Family |  |
| (Thorpe, 2024) | Isopoda | Family |  |
| (Copilaş-Ciocianu et al., 2020) | Amphipoda | Family |  |
| (Sun and Cheng, 2023) | Diplostraca | Family |  |
| (Eyun, 2017) | Copepoda | Family |  |
| (Yu et al., 2024) | Collembola | Order |  |
| (Misof et al., 2014) | Insecta | Order | All order-level constraints were followed except for Mecoptera, which was constrained as a polytomy with Siphonaptera to reflect recent evidence supporting the paraphyly of Mecoptera (Meusemann et al., 2020; Tihelka et al., 2020). |
| (Chang et al., 2020) | Orthoptera | Family |  |
| (Song et al., 2024) | Hemiptera | Family |  |
| (Blaimer et al., 2023) | Hymenoptera | Family | (Pteromalidae + Encytridae + Trichogrammatidae) and (Tenthredinidae + Athaliidae) were left as polytomies in the constraint tree due to non-monophyly. |
| (Zhang et al., 2018) | Coleoptera | Family |  |
| (Nie et al., 2020) | Chrysomelidae | Subfamily |  |
| (Kawahara et al., 2019) | Lepidoptera | Family |  |
| (Tihelka et al., 2020) | Siphonaptera + Mecoptera | Family |  |
| (Wiegmann et al., 2011) | Diptera | Family |  |
| (Suvorov et al., 2022) | Drosophilidae | Species |  |
| (daSilva et al., 2020) | Culicidae | Genus |  |

#### **Supplementary Table S3. Hypothesized transitions in germ cell specification in Panarthropoda.** Clade name = monophyletic group in whose ancestral lineage the transition is specified to have occurred; Node_number = node labels in Figures S1 and S2; Transition = gain of germ plasm (GP) or reversion to induction (IND); Parsimony = support from maximum parsimony (Figure S1); Likelihood = support from maximum likelihood (Figure S2).

See Table S3 at https://github.com/rishabhrajkapoor/panarthropoda_gc_specification_evolution, under supplementary_tables, commit ID ff9945c.

**Supplementary Table S4. Genomic and transcriptomic datasets used for phylogenetic inference.** We were unable to find sequence data for species in the “sub_species” columns, and instead substituted the genomes or transcriptomes for species in the same taxonomic family. Species in the “sub_species” columns are replaced by “iq_tree_label” in trees.

See Table S4 at https://github.com/rishabhrajkapoor/panarthropoda_gc_specification_evolution, under supplementary_tables, commit ID ff9945c.

#### **Supplementary Table S5. Literature-derived fossil calibrations.** Node = clade name calibrated by fossil bounds; Spanning set = list of two taxa spanning the full clade in our tree; Min = minimum node age (millions of years) used for soft fossil constraints in MCMCTree; Max = maximum node age used for soft fossil constraints in MCMCTree. In the Node column, “+” indicates a node representing the common ancestor of two monophyletic lineages. Note that Mecoptera was treated as paraphyletic in this study (see Table S2), such that the “Siphonaptera and Mecoptera” node represents the common ancestor of all Mecoptera and Siphonaptera.

| **Node** | **Spanning set** | **Min** | **Max** | **Reference** |
| --- | --- | --- | --- | --- |
| Panarthropoda | Parachela*, Drosophila_virilis* | 528.8 | 636.1 | (Howard et al., 2022) |
| Chelicerata | Nymphonidae*, Parasteatoda_tepidariorum* | 509.0 | 636.1 | (Howard et al., 2022) |
| Euchelicerata | Pseudoscorpiones*, Parasteatoda_tepidariorum* | 500.5 | 636.1 | (Howard et al., 2022) |
| Arachnopulmonata | Euscorpiidae*, Parasteatoda_tepidariorum* | 435.2 | 514.0 | (Howard et al., 2022) |
| Holometabola | Liposcelididae*, Camponotus_floridanus* | 306.9 | 450.0 | (Misof et al., 2014) |
| Pterygota | *Epiophlebia_superstes, Drosophila_virilis* | 324.0 | 450.0 | (Misof et al., 2014) |
| Dicondylia | *Thermobia_domestica, Drosophila_virilis* | 411.5 | 450.0 | (Misof et al., 2014) |
| Thysanoptera + Hemiptera | Phlaeothripidae*, Rhodnius_prolixus* | 207.0 | 450.0 | (Misof et al., 2014) |
| Hymenoptera | Tenthredinidae*, Camponotus_floridanus* | 221 | 450.0 | (Misof et al., 2014) |
| Formicidae + Apidae | *Camponotus_floridanus, Apis_mellifera* | 89.8 | 221.0 | (Misof et al., 2014) |
| Gyrinidae + Carabidae | *Gyrinus_marinus, Elaphrus_aureus* | 205.6 | 306.0 | (Misof et al., 2014) |
| Tenebrionidae + Curculionidae | *Sitophilus_oryzae, Tribolium_castaneum* | 207.0 | 306.0 | (Misof et al., 2014) |
| Lepidoptera | Hepialidae*, Bombyx_mori* | 112.0 | 306.9 | (Misof et al., 2014) |
| Papilionoidea | Papilionidae*, Pararge_aegeria* | 34.1 | 112.0 | (Misof et al., 2014) |
| Siphonaptera and Mecoptera | *Ctenocephalides_felis, Boreidae* | 160.5 | 306.9 | (Misof et al., 2014) |
| Diptera | *Phlebotomus_papatasi, Drosophila_virilis* | 230.0 | 306.9 | (Wiegmann et al., 2011) |
| Schizophora | *Bactrocera_tryoni, Drosophila_virilis* | 64.0 | 230.0 | (Wiegmann et al., 2011) |
| Sciaroidea | *Bradysia_coprophila, Mayetiola_destructor* | 180.0 | 230.0 | (Wiegmann et al., 2011) |
| Arthropoda | *Limulus_polyphemus, Drosophila_virilis* | 514.0 | 636.1 | (Bernot et al., 2023) |
| Branchiopoda | Triopsidae*, Daphnia_magna* | 405.0 | 521.0 | (Bernot et al., 2023) |
| Collembola | Tomoceridae*, Tetrodontophora_bielanensis* | 405.0 | 521.0 | (Bernot et al., 2023) |
| Peracarida | Gammaridae*, Armadillidium_nasatum* | 358.2 | 521.0 | (Bernot et al., 2023) |

**Supplementary Table S6. phytools model fits for maximum likelihood ancestral state reconstruction.** SYM = symmetric substitution rates; ARD = asymmetric substitution rates; SYM.G = symmetric substitution rates with gamma heterogeneity; ARD.G = asymmetric substitution rates with gamma heterogeneity; HRM2 = hidden rates model with symmetric substitution rates and two hidden rate categories. log(L) = log likelihood score; d.f. = degrees of freedom (number of model parameters); AIC = Akaike information criterion; Weight = model importance via ANOVA.

| **Model** | **log(L)** | **d.f.** | **AIC** | **Weight** |
| --- | --- | --- | --- | --- |
| SYM | -68.10199 | 1 | 138.204 | 0.0065576866 |
| ARD | -67.93943 | 2 | 139.879 | 0.0028382863 |
| SYM.G | -62.41206 | 2 | 128.824 | 0.7137752031 |
| ARD.G | -62.36246 | 3 | 130.725 | 0.2759361780 |
| HRM2 | -67.09620 | 4 | 142.192 | 0.0008926461 |

#### **Supplementary Table S7. Marginal probabilities at select nodes.** Probabilities of germ plasm nodes at select ancestors derived via maximum likelihood. Node_number = labels at internal nodes of Figures S1 and S2; IND = induction; GP = germ plasm.

| **Clade** | **Node_number** | **P_IND** | **P_GP** |
| --- | --- | --- | --- |
| Panarthropoda | 1 | 0.994 | 0.006 |
| Arthropoda | 2 | 1.000 | 0.000 |
| Chelicerata | 3 | 1.000 | 0.000 |
| Myriapoda | 4 | 0.994 | 0.006 |
| Pancrustacea | 5 | 1.000 | 0.000 |
| Decapoda | 6 | 0.119 | 0.881 |
| Hyalidae + Talitridae | 7 | 0.084 | 0.916 |
| Poduromorpha + Symphypleona | 8 | 0.668 | 0.332 |
| Collembola | 9 | 0.987 | 0.013 |
| Insecta | 10 | 0.999 | 0.001 |
| Holometabola | 11 | 0.806 | 0.194 |
| Hymenoptera | 12 | 0.283 | 0.717 |
| Coleoptera | 13 | 0.986 | 0.014 |
| Chrysomelidae + Curculionidae | 14 | 0.120 | 0.880 |
| Lepidoptera | 15 | 0.996 | 0.004 |
| Mecoptera + Siphonaptera | 16 | 0.518 | 0.482 |
| Antliophora (Diptera + Mecoptera + Siphonaptera) | 17 | 0.532 | 0.468 |
| Diptera | 18 | 0.385 | 0.615 |

**Supplementary Table S8. Results of Pagel’s test of correlated rates in germ band type and germ cell specification mode.**

| **Model** | **log(L)** | **AIC** | **p-value** |
| --- | --- | --- | --- |
| Independent | -79.52302 | 167.0460 | n/a |
| Bidirectional | -65.03560 | 146.0712 | 7.90998e-06 |
| GC dependent | -69.61696 | 151.2339 | 4.98718e-05 |
| GB dependent | -68.75534 | 149.5107 | 2.10696e-05 |

**Supplementary Table S9. Results of search for *oskar* orthologs.** BUSCO scores may differ from Table S4 for the seven unannotated genomes used in our analysis (*Clogmia albipunctata, Mayetiola destructor, Macrocentus cingulum, Epiphyas postvittana, Nymphalis antiopa, Pyrrhocoris apterus, Tribolium confusum*), as BUSCO scores and *oskar* presence are determined from AUGUSTUS genome annotations. “Blondel_oskar_count” column indicates the total number of *oskar* orthologs previously identified across all datasets from this species (Blondel et al., 2021). Empty cells indicate species that were not analyzed in our previous study on *oskar* ortholog evolution (Blondel et al., 2021). “oskar_present” column indicates that *oskar* was detected in the current analysis and/or in our previous work (Blondel et al., 2021). “data_type” indicates whether the protein predictions were obtained directly from protein fasta files on NCBI (“NCBI-annotated genome”), from TransDecoder run on transcriptome shotgun assembly nucleotide sequences (“TSA”), or from *ab initio* annotations generated with AUGUSTUS (“AUGUSTUS-annotated genome”). We were unable to find sequence data for species in the “sub_species” column, and instead substituted the genomes/transcriptomes for species in the same taxonomic family. The “oskar_accession” columns provide either NCBI protein accessions for NCBI-annotated genomes (“https://www.ncbi.nlm.nih.gov/protein/”), TSA nucleotide accessions for TSAs (“https://www.ncbi.nlm.nih.gov/genbank/tsa/”), or AUGUSTUS-generated accessions for AUGUSTUS-annotated genomes. All Oskar protein sequences are available in oskar_proteins.fa.

See Table S9 at https://github.com/rishabhrajkapoor/panarthropoda_gc_specification_evolution, under supplementary tables, commit ID ff9945c.
